# miR-155-5p/miR-674-3p presence in peripheral blood leukocytes and relative proportion of white blood cell types as biomarkers of asymptomatic and symptomatic phases of temporal lobe epilepsy

**DOI:** 10.1101/2024.02.29.582734

**Authors:** Kinga Szydłowska, Piotr Chrościcki, Maciej Olszewski, Karolina Nizińska, Katarzyna Piwocka, Katarzyna Łukasiuk

## Abstract

Epilepsy frequently develops as a result of brain insult, for example, brain injury or stroke. Currently, there are no tools allowing us to predict which trauma patients will eventually develop epilepsy. There is evidence that microRNAs levels are altered in the blood, making them attractive candidates for peripheral biomarkers of epilepsy. We analyzed white blood cell subpopulations containing miR-155-5p and miR-674-3p, in control and stimulated animals and in control and symptomatic or asymptomatic animals in the amygdala stimulation model. The first proposed early biomarker of epilepsy is the relative proportion of CD45RA^+^ B cells containing miR-155-5p and/or miR-674-3p. Others are increased number of CD45RA^+^ B cells containing either miR-155-5p or miR-155-5p and miR-674-3p together or decreased number of CD161^+^ NK cells not containing miR-155-5p nor miR-674-3p. Additionally, we found that the decreased number of CD4^+^ T cells can be used as a potential biomarker for identifying epileptic animals with symptomatic epilepsy.

## Introduction

Epilepsy is one of the most common neurological disorders responsible for severe disabilities in 65 million of patients worldwide (Milligan, 2021). One of the most common causes of epilepsy in adults are traumatic brain injury (TBI) and stroke. The incidence of epilepsy in patients who suffered from TBI is 25% at five years and 32% at 15 years (Pease *et al*, 2022). Among patients with one late seizure > 7 days post-trauma, the risk of seizure recurrence was 62% after one year and 82% at ten years years (Pease *et al*., 2022). The incidence of early post-stroke seizures (occurring ≤ one week from stroke onset) and late post-stroke seizures (occurring > one week from stroke onset) is 4.5% and 2.1%, respectively (Lee *et al*, 2022). Due to the lack of diagnostic tools, there is no possibility of predicting which patients should be monitored more closely or require an introduction of early treatment due to the high risk of developing epilepsy which makes testing of potential antiepileptogenic therapies much more difficult.

Pre-clinical studies are indispensable for drug studies or the discovery of basic mechanisms of diseases. In the case of epilepsy studies, identification of animals suffering from epilepsy requires time and labor-consuming analysis of EEG recordings in search for electrographic seizures (Engel *et al*, 2013). Thus there is a dire need for the identification of easily tested and precise biomarkers that would allow early prediction of epilepsy development or allow assessing the effect of antiepileptogenic or antiepileptic therapeutics.

MicroRNAs could be good candidates for biomarkers of epileptogenesis in patients and experimental animals or epilepsy phenotype in preclinical studies (De Benedittis *et al*, 2021; Henshall, 2013; Henshall *et al*, 2016; Jimenez-Mateos & Henshall, 2013; Pitkänen *et al*, 2019; Pitkänen & Lukasiuk, 2011; Simonato, 2018; Simonato *et al*, 2021). MicroRNAs are 19-22 nucleotide long RNA molecules that can regulate the target mRNA expression (Rao *et al*, 2013). MicroRNAs come from endogenous transcripts that form hairpin structures, which are processed such that a single miRNA molecule accumulates from one arm of a hairpin precursor molecule (Ambros *et al*, 2003). It is estimated that a single miRNA can regulate up to hundreds of genes, which makes them one of the most powerful regulators of gene expression (Ebert & Sharp, 2012).

miRNAs are widely expressed in the brain, both in neuronal and glial cells (Bot *et al*, 2013; Jovičić *et al*, 2013; Sempere *et al*, 2004). Alterations in levels of miRNA in the brain in epilepsy and epileptogenesis have been shown in experimental animals and human epilepsy (Bot *et al*., 2013; Gorter *et al*, 2014; Hu *et al*, 2011; Roncon *et al*, 2015; Srivastava *et al*, 2017; Zucchini *et al*, 2014).

Interestingly, changes in levels of miRNAs associated with neurological diseases have been shown also in body fluids, including plasma and blood (Engel *et al*., 2013; Lukasiuk & Becker, 2014; Pitkänen *et al*, 2018; Venereau *et al*, 2016; Walker *et al*, 2017; Walker *et al*, 2016). In epilepsy, changes in miRNAs levels have been found in the plasma of patients with focal resistant epilepsy (Pollard *et al*, 2012), serum of patients with epilepsy (Wang *et al*, 2015a; Wang *et al*, 2015b), plasma of mice in pilocarpine model (Roncon *et al*., 2015), plasma of mice in pilocarpine model, perforant path stimulation model in rats, and patients with focal epilepsy (Brennan *et al*, 2020), in plasma of patients with refractory focal epilepsy (Raoof *et al*, 2018), whole blood from rats following kainate-induced seizures (Liu *et al*, 2010), in the blood of mice following pilocarpine-induced status epilepticus (Hu *et al*., 2011) and in the plasma of rats with amygdala stimulation induced epilepsy (Szydlowska *et al*, 2024).

Although whole blood has been studied in the context of miRNA changes in epilepsy, there is no data on the cellular localization of candidate miRNAs despite the interplay between the immune system and the brain, being a highly multilayered and well-proven interaction (Marchi *et al*, 2014; Steinman, 2004; Vezzani *et al*, 2011; Vezzani & Granata, 2005). It was shown that the immune cells may travel between the brain and the blood, also some immune cells from the blood may infiltrate the brain, especially in a diseased state (Yamanaka *et al*, 2021). Therefore, full blood may be a better source of biomarkers than plasma or serum.

In this study, we aimed at a multimethod search of early biomarkers of epileptogenesis and epilepsy in the blood in a well-characterized model of epilepsy induced by status epilepticus evoked by the amygdala stimulation in rats (Nissinen *et al*, 2000; Nizinska *et al*, 2021). In this model, epileptogenesis-triggering status epilepticus is followed by a long, seizure-free latency period and then the appearance of spontaneous, unprovoked seizures. We conducted experiments to detect miRNA differentiating between different phases and phenotypes of epilepsy development, and to define the cellular localization of selected hits.

This is a proof of concept study showing the possibility of using blood miRNA and cellular localization of individual miRNAs in white blood cell populations to predict epilepsy development and differentiation between epileptic vs. nonepileptic injured individuals. This is also the first study, which presents the analysis of miRNAs levels in different white blood cell subpopulations during epileptogenesis and epilepsy.

## Results

### Status epilepticus evoked by the amygdala stimulation alters miRNA levels in the blood

We identified alterations in levels of 56 miRNAs at 2 days, 66 miRNAs at 7 days, and 19 miRNAs at 8 months after stimulation (p<0.05) (Fig 1A). 3D PCA analysis revealed clear spatial separation of samples from control and stimulated animals at each tested time point (Fig 1B). Additionally, the separation between early (2 and 7 days) and late (8 months) time points was detected. (Fig 1C).

**Figure 1.**
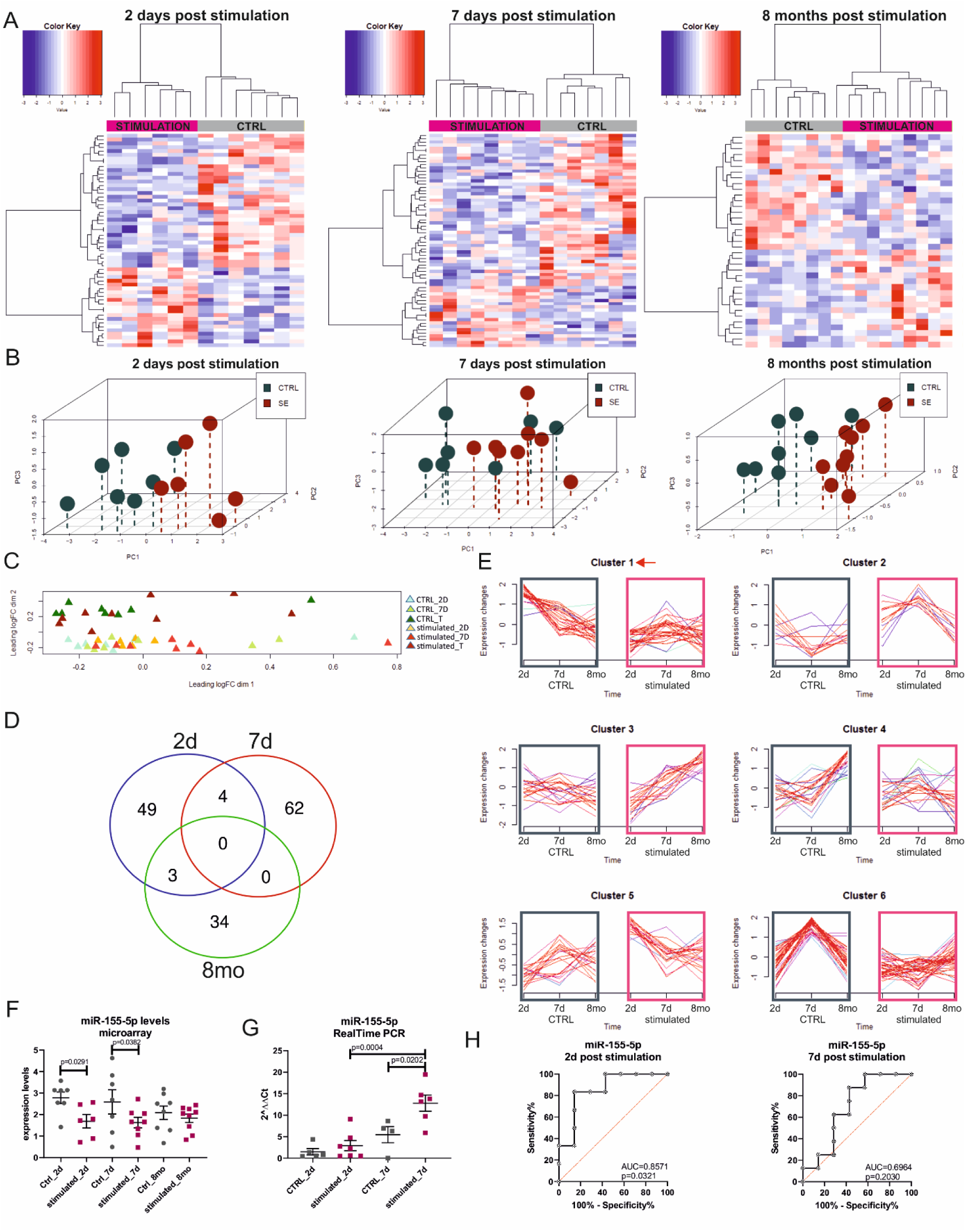
Amygdala stimulation affects miRNA levels. A Heatmaps showing miRNAs signatures differentiating between control and stimulated animals at different time points. B Three dimensional PCA graphs present spatial separation of samples from control and stimulated animals for each time point tested based on miRNAs signatures differentiating between control and stimulated animals. Gray bars – control animals, pink bars – stimulated animals. C Three dimensional PCA graph of animal samples from 2 days, 7 days and 8 months after stimulation based on miRNAs signatures differentiating between control and stimulated animals at each time point. D Venn diagram presenting numbers of miRNAs significantly changing level at each time point. E Supervised clustering of miRNA level over time. Red arrow points to the cluster 1 containing miR-155-5p. Gray squares – control animals, pink squares – stimulated animals. F Dot plot of miR-155-5p level detected by microarrays in discovery cohort of animals. G Dot plot of miR-155-5p level detected using Real Time PCR in validation cohort of animals. Analyzed using ΔΔCt method. H ROC analysis presenting percentage of sensitivity and specificity for difference in the level of miR-155-5p between control and stimulated animals, at 2 and 7 days after stimulation. Data information: For (A-E) data analyzed using R. For the expression analysis, the calculated p-values were based on moderated t-statistics. Furthermore, the Benjamini and Hochberg multiple testing adjustment method was applied to the p-values (FDR – False Discovery Rate). A one-way ANOVA was used to establish miRNA with differential levels between the tested groups. Controls are time-matched for each time point tested. For (F-G) data are shown as mean ± SEM. P values are determined by two-way ANOVA. All presented data come from 2 cohorts of animals combined together (total n was 11 control and 15 stimulated animals). Source data are available online for this figure. Controls are time-matched for each time point tested.

There are 4 miRNAs (miR-155-5p, miR-632, miR-150-5p, and miR-200c-3p) which levels change significantly between control and stimulated animals at both 2 and 7 days post-stimulation (Fig 1D). No alterations for these miRNAs were found at 8 months after stimulation (Fig 1D). miRNA miR-155-5p was chosen as a primary candidate for a biomarker for further studies.

Supervised clustering of miRNAs levels changes over time into 6 elements’ clusters revealed different patterns of expression dynamics between control and stimulated animals. (Fig 1E). Three clusters contained miRNAs fluctuating in control but not in stimulated animals (Fig 1E – clusters 1,4, and 6). Cluster 3 grouped miRNAs with stable levels over time in control animals and increasing levels in stimulated animals (Fig 1E). In cluster 2, levels of miRNAs at 7 days post-stimulation decreased in control animals and increased in stimulated animals (Fig 1E).

miR-155-5p, which was selected as a primary candidate for biomarker, was assigned to cluster 1 (Fig 1E). In control animals, miRNAs from this cluster strongly increased in level at 2 days, and 7 days, and then returned to their basic level at 8 months. Interestingly, in stimulated animals, the level of miR-155-5p remained low at all tested time points (stimulated animals: 2d = 1.69, 7d = 1.63, 8mo = 1.83; Controls: 2d = 2.78, 7d = 2.59, 8mo = 2.08; 2d p = 0.0291, 7d p = 0.0382). Alterations in levels of miR-155-5p were validated by Real-Time PCR using blood samples from a new cohort of animals. This analysis confirmed our finding that miR-155-5p levels change (Fig 1 F-G). The level of miR-155-5p in stimulated animals at 2 and 7 days was lower than in control animals. ROC analysis of microarray data revealed high sensitivity and specificity of miRNA-155-5p differentiation between control vs stimulated animals at 2 days after stimulation (2d AUC = 0.8571; Fig 1H).

### miRNAs signatures in epileptic animals differ, depending on the duration of the latency phase

Long-term analysis of EEG performed 24 hours a day, 7 days a week, for 3 months in total, revealed that 4 of the stimulated animals developed epilepsy (that is experienced first spontaneous seizure) by the 8^th^ day (an animal group referred to as: EARLY) and 4 animals had first spontaneous seizures later than 20 days post-stimulation (an animal group referred to as LATE) (Fig 2E).

**Figure 2.**
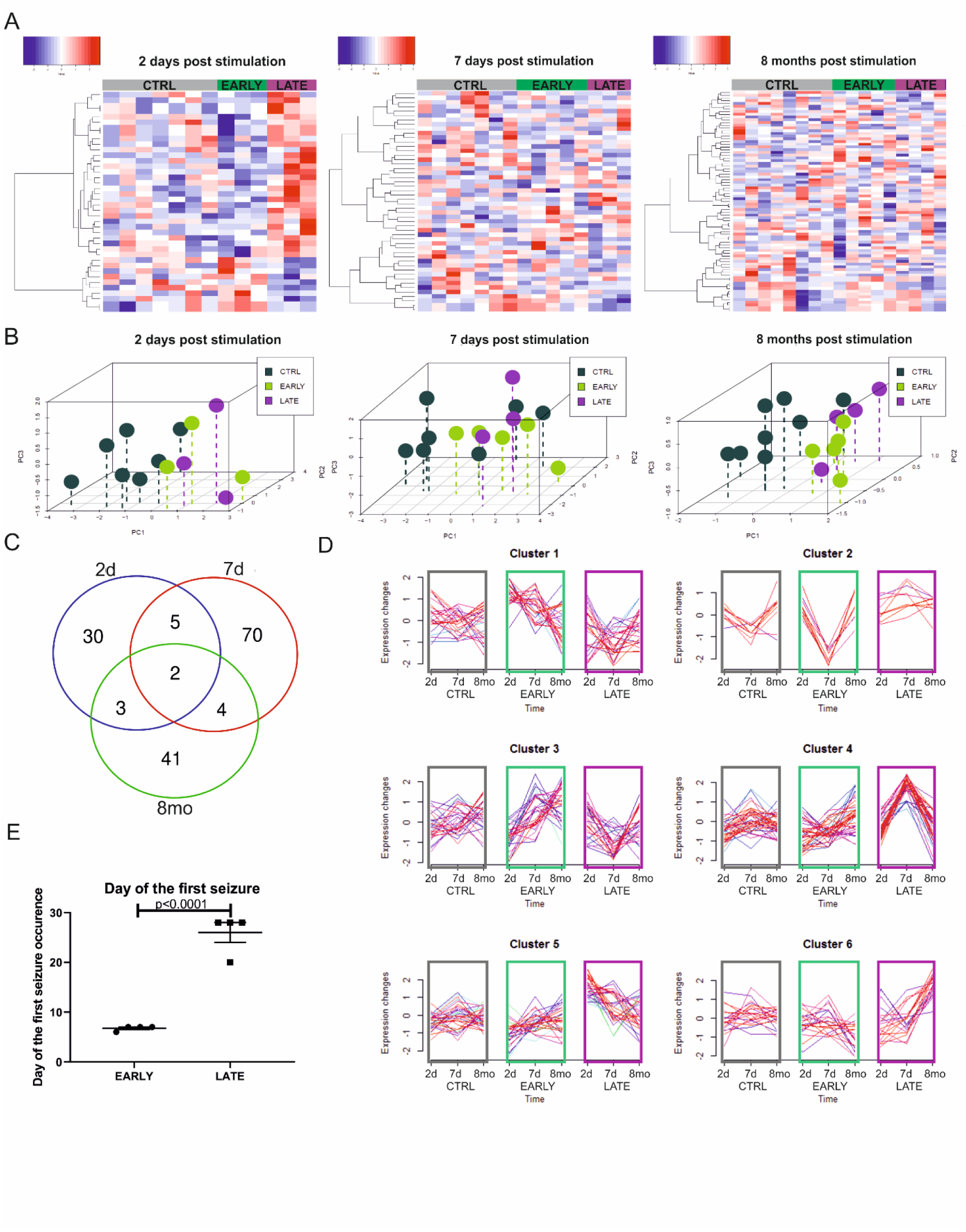
Early or late seizures occurrence in stimulated animals affects miRNA levels. For (A-E) data, EARLY group refers to a group of rats which experienced first seizure at day 6-7 post stimulation and LATE group refers to animals that experienced first seizure later than 20 days post stimulation. A Heatmaps show selected miRNAs, significantly changing levels between EARLY and LATE animals. B Three dimensional PCA graphs present special separation of samples from control, EARLY and LATE animals for each time point tested based on miRNAs with significant difference between EARLY and LATE animals. Gray bars – control animals, green bars – EARLY animals and purple bars – LATE animals. C Venn diagram presenting number of miRNAs significantly changing between EARLY and LATE animals in each time. D Supervised clustering of miRNA levels in control, EARLY and LATE animals over time. Gray squares – control animals, green squares – EARLY animals and purple squares – LATE animals. E Dot plot presenting day of the first seizure experienced by animals. Data information: For (A-D) data analyzed using R. For the expression analysis, the calculated p-values were based on moderated t-statistics. Furthermore, the Benjamini and Hochberg multiple testing adjustment method was applied to the p-values (FDR – False Discovery Rate). A one-way ANOVA was used to establish miRNA with differential levels between the tested groups. For (E) data are shown as mean ± SEM. P value is determined by t test. All presented data come from 2 discovery cohorts of animals combined together (total n was 11 control and 15 stimulated animals). Source data are available online for this figure. Controls are time-matched for each time point tested.

When comparing EARLY and LATE animal groups, we detected 42 miRNAs with levels differentiating between groups at 2 days post-stimulation, 50 miRNAs at 7 days post-stimulation, and 81 miRNAs at 8 months post-stimulation (Fig 2A). Two miRNAs, miR-31a and miR-31b, differentiated significantly between EARLY and LATE animal groups at all tested time points (Fig. 2C). There were 5 miRNAs with levels differentiating at 2 and 7 days after stimulation, and 3 miRNAs at 2 days and 8 months’ time points. Finally, 4 miRNAs changed their levels significantly between 7 days and 8 months samples (Fig. 2C).

Differences in signatures of miRNAs between EARLY and LATE groups were detected in all time points; however, the most pronounced differences were observed at 2 days after stimulation (Fig. 2A). Supervised clustering confirmed that at 2 days post-stimulation the samples from EARLY (green bars on Fig 2B) and LATE (purple bars on Fig 2B) animals were separated.

Supervised clustering of miRNAs differentiating between EARLY and LATE groups, identified clusters of miRNAs with similar changes of their level patterns over time. Two clusters contained miRNAs fluctuating with time in EARLY epileptic animals, whereas miRNAs from LATE animals were stable over time (Fig 2D – clusters 1 and 2). Cluster 3 grouped miRNAs with signatures increasing over time in EARLY and decreasing over time in LATE epileptic animals (Fig 2D). Remaining clusters 4, 5, and 6, grouped mRNAs whose levels were stable in EARLY epileptic animals and increased post-stimulation in LATE epileptic animals: at 2 days in cluster 5, at 7 days in cluster 4 and at 8 months in miRNAs from cluster 6 (Fig 2D).

### Animals with low vs high numbers of seizures differ in miRNAs signatures

While analyzing EEG data, we identified two subgroups of rats in the stimulated animals group, on the basis of the number of seizures during the observation period (3 months in total) (Fig 3E). The average seizure number was 210.7 ± 254.2. We assigned animals with lower than average (60.8 ± 68.8) seizures number to the LOW group and with higher than average (460.7 ± 258.4) seizures number to the HIGH group.

**Figure 3.**
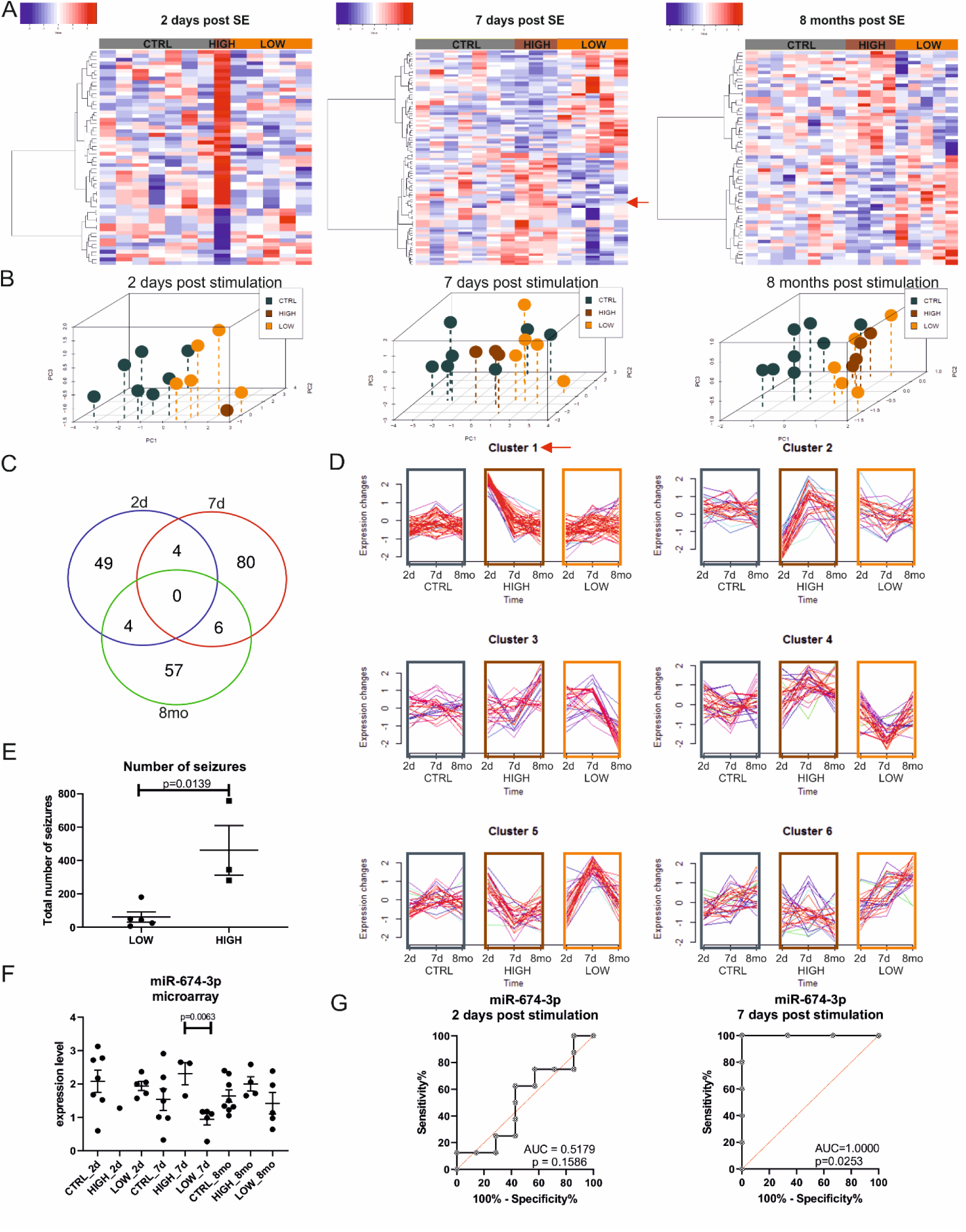
Epilepsy intensity, calculated as seizure number, induces changes of miRNA levels in stimulated animals. For (A-G) data, HIGH group refers to a group of rats with high total number of seizures experienced post stimulation and LOW group refers to animals with low total number of seizures experienced post stimulation. A Heatmaps show levels of selected miRNAs, significantly changing between HIGH and LOW animals. B Three dimensional PCA graphs present special separation of samples from control and stimulated animals for each time point tested. Based on selected miRNAs with significant difference between HIGH and LOW animals. Gray bars – control animals, brown bars – HIGH animals and orange bars – LOW animals. C Venn diagram presenting number of miRNAs significantly changing between HIGH and LOW animals in each time point separately and in more than one time point. D Levels of selected miRNAs clustered using Mfuzz algorithm in R, to identify groups of miRNAs with levels changing in a similar way over time between animals with HIGH and LOW number of seizures. Red arrow points to the cluster containing selected miR-674-3p. Gray squares – control animals, brown squares – HIGH animals and orange squares – LOW animals. E Dot plot presenting number of seizures for HIGH and LOW group of animals. Groups were split based on the mean of seizure number for all the animals. F Dot plot of miR-674-3p level detected by microarrays in discovery cohort animals. G ROC analysis presenting percentage of sensitivity and specificity for difference in level of miR-674-3p between HIGH and LOW animals, at 2 and 7 days after stimulation. Data information: For (A-E) data analyzed using R. For the expression analysis, the calculated p-values were based on moderated t-statistics. Furthermore, the Benjamini and Hochberg multiple testing adjustment method was applied to the p-values (FDR – False Discovery Rate). A one-way ANOVA was used to establish miRNA with differential levels between the tested groups. Controls are time-matched for each time point tested. For (F-G) data are shown as mean ± SEM. P values are determined by two-way ANOVA. All presented data come from 2 cohorts of animals combined together (total n was 11 control and 15 stimulated animals). Source data are available online for this figure. Controls are time-matched for each time point tested.

We identified 57 miRNAs with levels differentiating between HIGH and LOW groups at 2 days, 90 at 7 days, and 67 miRNAs at 8 months post-stimulation (Fig 3A). 3D PCA analysis showed strong spatial separation between the animals with HIGH (brown bars in Fig 3B) and LOW (orange bars in Fig 3B) numbers of seizures. Particularly clear differences in the separation of animals samples were obtained for rats with HIGH and LOW numbers of seizures at 7 days post stimulation (Fig 3B). At 8 months post stimulation, differentiation between the animals with HIGH and LOW numbers of seizures was still detected (Fig 3B).

Signatures of six miRNAs differentiated between HIGH and LOW at 2 days, 7 days and 8 months after stimulation (Fig 3C). As a candidate for a biomarker for further studies, we selected miR-674-3p, which had a high level and also high Fold Change (Fig 3F) and maximum sensitivity and specificity in differentiating between animals with HIGH vs LOW seizures number at 7 days after stimulation (AUC=1.000) (Fig 3G). The level of miR-674-3p did not differ between HIGH vs. LOW groups at 8 months (Fig 3F).

Unsupervised clustering into 6 clusters of miRNAs level changes over time for control and HIGH and LOW groups of animals revealed several patterns (Fig 3D). Some miRNAs have stable levels over time in animals with a LOW number of seizures, but the levels of those miRNAs in animals with the HIGH number of seizures increased (cluster 1) or decreased (cluster 2) at 2 days after stimulation (Fig 3D). Other interesting clusters are clusters 4, and 5, where observed changes of miRNA levels in LOW and HIGH groups were opposite (Fig 3D). miR-674-3p was assigned to cluster 1 in which, in control animals and animals with the low number of seizures, miRNAs levels were low and stable over time. In contrast, in animals with the high number of seizures, its level increased at 2 and 7 days post-stimulation (that is, before epilepsy diagnosis) and returned to the control level at later time points (Fig 3D and F). ROC analysis revealed high sensitivity and specificity of change in miR-674-3p level at 2 days (AUC = 0.5179) and at seven days (AUC = 1.0000) post-stimulation (Fig 3H).

### Numbers of CD4^+^ T cells decrease in the blood of epileptic animals

Basic blood morphology analysis performed on blood samples from the discovery group of animals revealed no differences in the number of erythrocytes, leukocytes, neutrophils, lymphocytes, monocytes, eosinophils, and basophils, between control and stimulated animals at 8 months after stimulation (Suppl Fig 1).

To characterize subpopulations of blood cells, we employed flow cytometry with fluorescent in situ RNA hybridization method (PrimeFlow RNA assay) and studied early and late time points after the stimulation (Fig 4A). For this experiment, a new validation cohort of animals was employed (n=6 stimulated and n=6 sham-operated rats). No significant differences were detected between control and stimulated animals at any time in numbers of CD8^+^ T cells, CD161^+^ NK cells, and CD45RA^+^ B cells (Suppl Fig 2). A significant decrease in stimulated animals was observed at 2 days post stimulation in the number of CD43^high^ monocytes (38.16% ± 8.36) and an increase in stimulated animals in the number of CD43^high^ granulocytes (182.93% ± 36.79) compared to control (Fig 4B and Suppl Fig 2). At 7 days post-stimulation there was an increase in the number of CD43^low^ granulocytes, and at 8 months post stimulation decrease in the number of CD4^+^ T cells in stimulated animals (Fig 4B and Suppl Fig 2).

**Figure 4.**
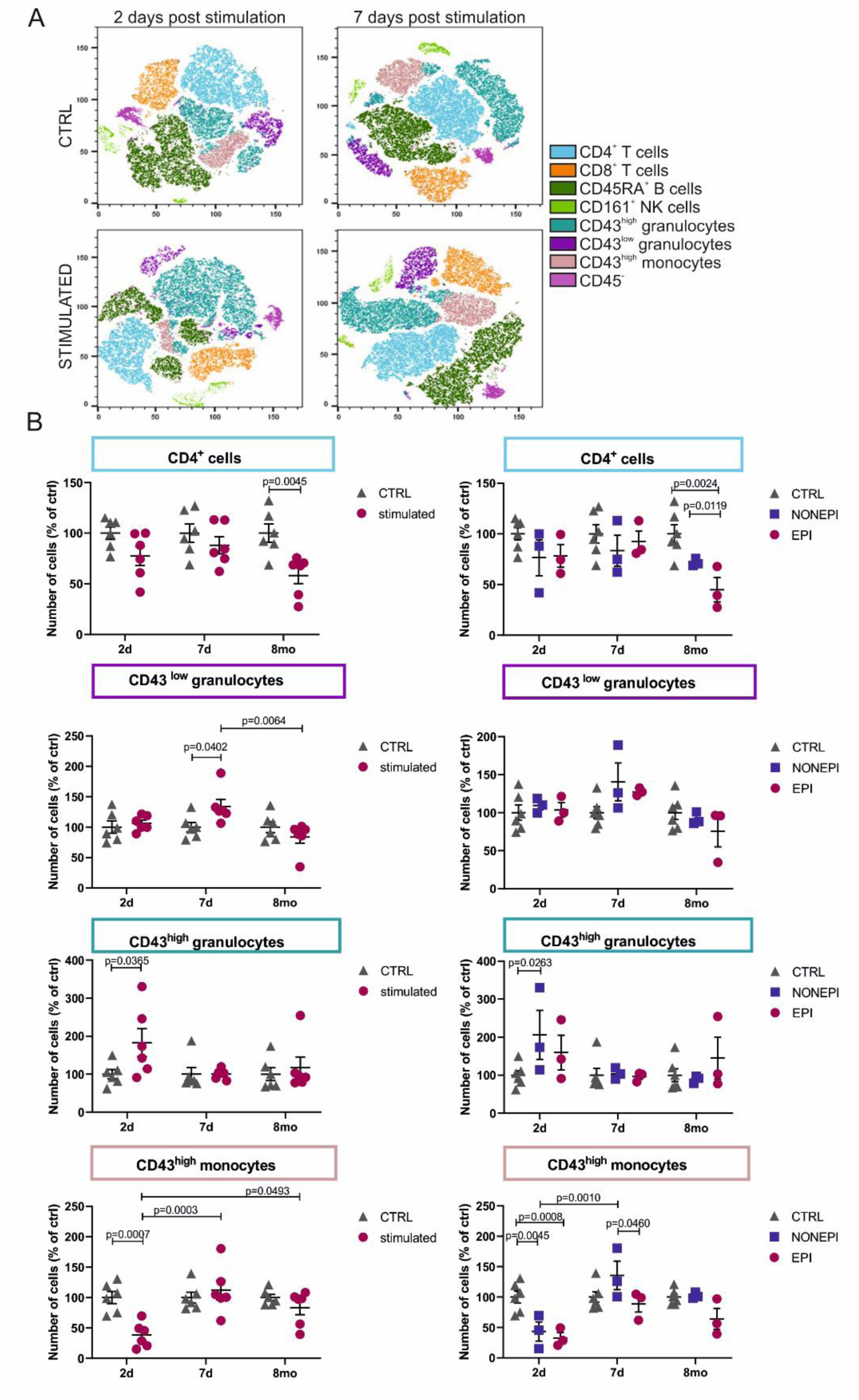
Effect of stimulation on leukocyte subpopulations. A tSNE plots present the cumulative 7 leukocytes subpopulations in control and stimulated animal blood samples at 2 and 7 days after stimulation. B Dot plots show percentage of cells in control and in stimulated animals or in animals that did (EPI) or did not (NONEPI) develop epilepsy. Graphs are shown only for cell types in which significant changes between any of the groups were detected. Data information: Data are shown as number of cells and percentage of control using time-matched controls. Data are shown as mean ± SEM. P value is determined by two-way ANOVA and Tukey’s multiple comparison test. All presented data come from one cohort of animals (Ctrl n = 6, stimulated n = 6, EPI n = 3, NONEPI n = 3).

In this cohort of animals, out of 6 stimulated animals, 3 did develop epilepsy (EPI group – symptomatic animals), and 3 did not (NONEPI group – asymptomatic group) during 8 months of the experiment. When comparing symptomatic and asymptomatic animal cell numbers, there was no difference detected at any time point in the number of: CD8^+^ T cells, CD161^+^ NK cells, and CD45RA^+^ B cells (Suppl Fig 2).

Interestingly, several differences in other types of white blood cell numbers were observed at all tested time points when comparing symptomatic (EPI) and asymptomatic (NONEPI) animals. At 2 days post-stimulation, there was an increase in the number of CD43^high^ granulocytes in asymptomatic animals and a decreases in the number of CD43^high^ monocytes both in asymptomatic and symptomatic animals when compared to control (Fig 4B). Particularly interesting was the decreased number of CD4^+^ T cells in symptomatic animals (EPI group) compared to both control and asymptomatic animals at 8 months post-stimulation (Fig 4B). Thus, we propose the number of CD4^+^ T cells as the biomarker of symptomatic epilepsy.

### The number of white blood cells subpopulations containing miR-155-5p and/or miR-674-3p changed after the stimulation

The use of the PrimeFlow method allowed co-detection of 2 selected miRNAs (miR-155-5p and miR-674-3p) in white blood cells subpopulations: CD4^+^ T cells, CD8^+^ T cells, CD161^+^ NK cells, CD45RA^+^ B cells, CD43^high^ granulocytes, CD43^low^ granulocytes, and CD43^high^ monocytes. The distribution of each subpopulation numbers in control and stimulated animals was presented as paired with heat dot plots representing the presence of each detected miRNA in Fig 5 (Fig 5). This analysis gives an overview of changes in miR-155-5p and miR-674-3p levels in different white blood cell subpopulations, between control and stimulated animals post-stimulation (Fig 5).

**Figure 5.**
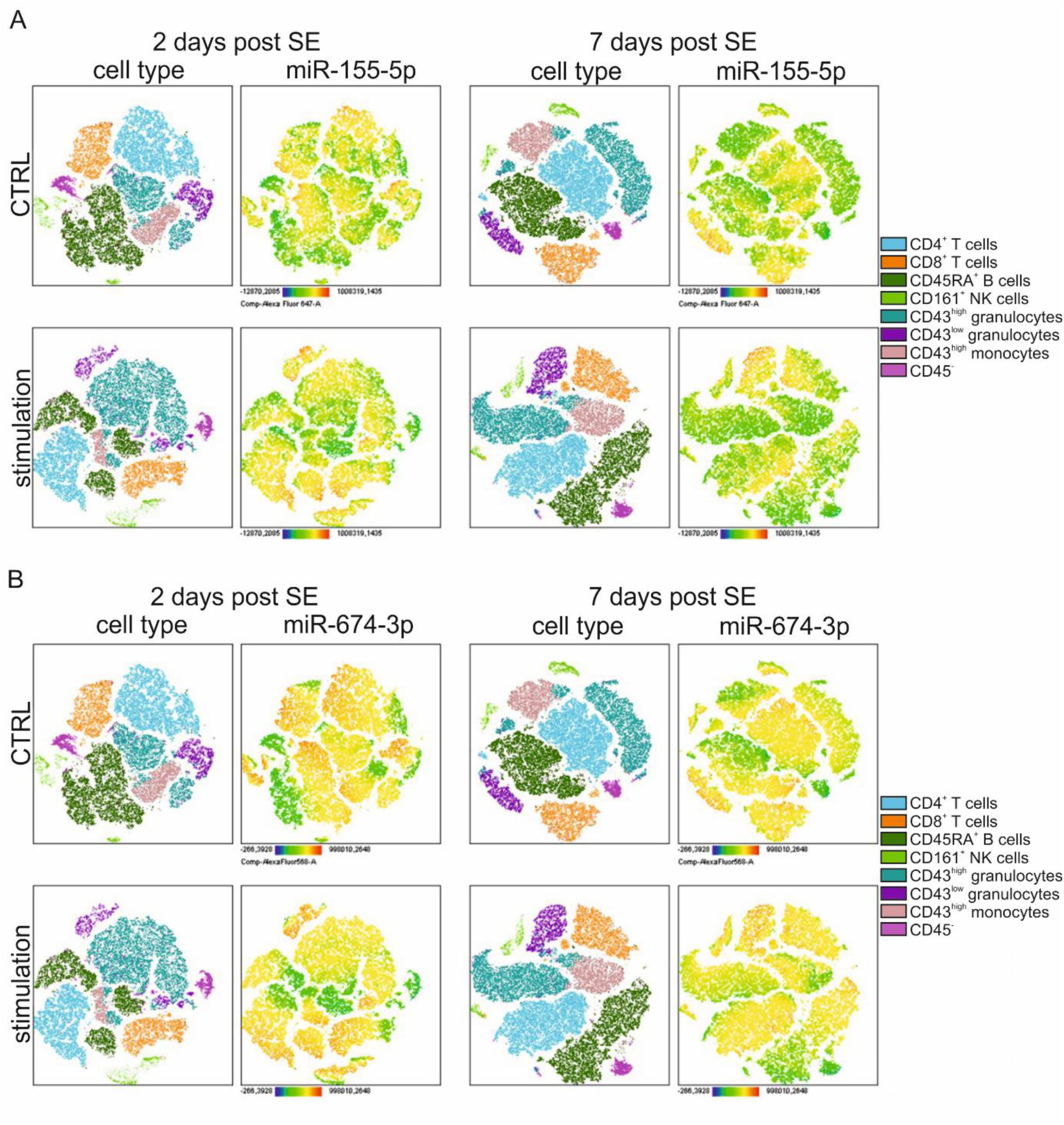
miR-155-5p and miR-674-3p are differentially distributed in various leukocytes subpopulations depending on stimulation status. A-B tSNE plots on the left side of each panel present seven white blood cells subpopulations. Heatmaps on the tSNE plots on the right of each panel present fluorescence intensity of intracellular miR-155-5p (panel A) and miR-674-3p (panel B) at 2 and 7 days post stimulation. Color code indicates low (blue) or high (red) content of miRNA in the cell.

We analyzed miRNA presence for both of the detected miRNAs, presenting a relative proportion of cells containing selected candidates for biomarkers (Fig 6). We evaluated 6 cell subclasses characterized by: (i) presence of miR-155-5p (could be co-stained with miR-674-3p), (ii) only miR-155-5p (no co-staining with miR-674-3p), (iii) presence of miR-674-3p (could be co-stained with miR-155-5p), (iv) presence of only miR-674-3p (no co-staining with miR-155-5p), (v) presence of both miR-155-5p and miR-674-3p and also (vi) lack of both miRNAs (Fig 6, Fig 7 and Fig 8).

**Figure 6.**
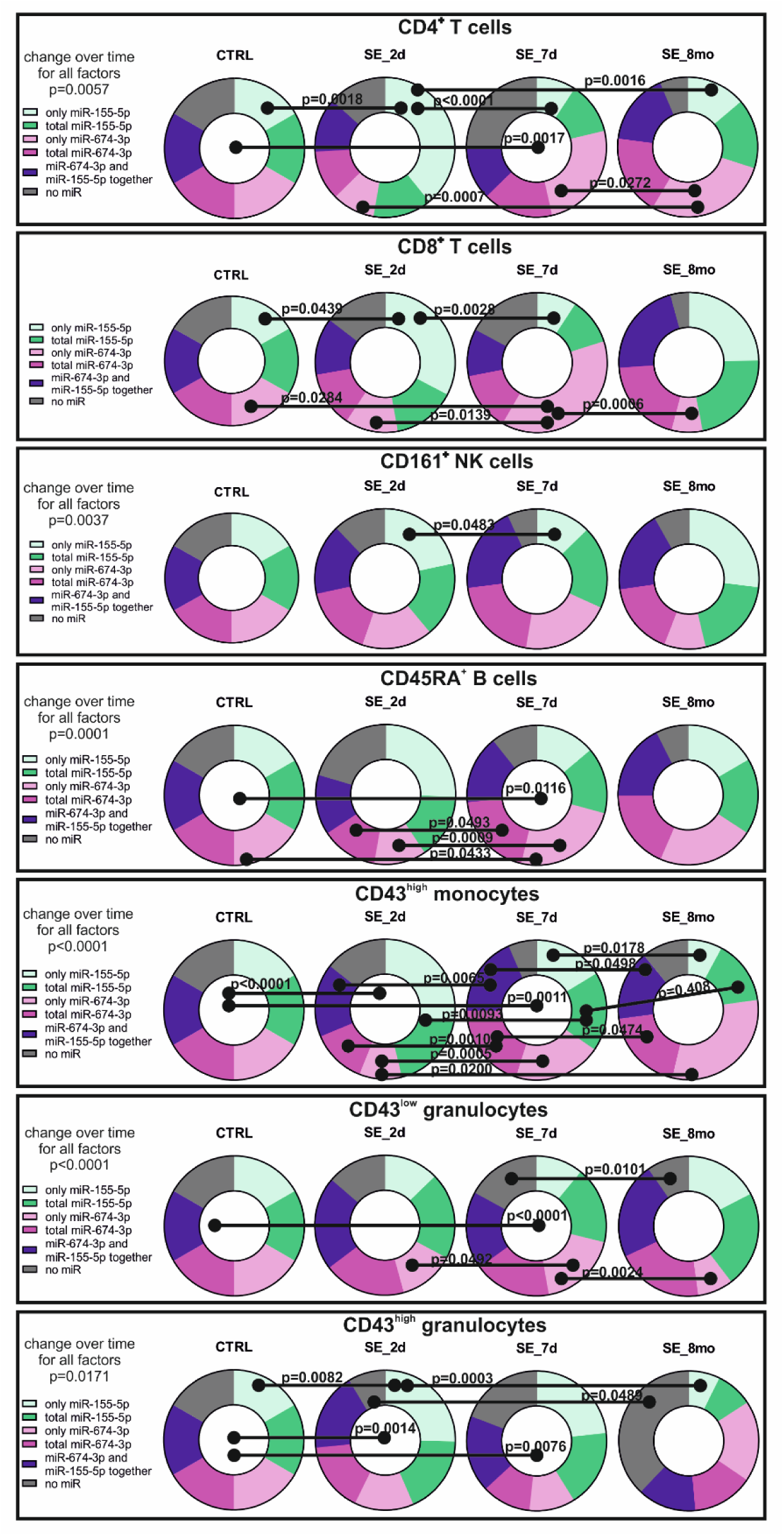
6 subclasses of white blood cells containing miR-155-5p and/or miR-674-3p differ between control and stimulated animals. Pie charts of relative proportion of white blood cells subpopulations containing miR-155-5p and/or miR-674-3p for control and stimulated animals at 2 days, 7 days and 8 months after stimulation. 6 subclasses of cells containing miR-155-5p and/or miR-674-3p: only miR-155-5p (no co-staining with miR-674-3p), total miR-155-5p (could be co-stained with miR-674-3p), only miR-674-3p (no co-staining with miR-155-5p), total miR-674-3p (could be co-stained with miR-155-5p), both miR-155-5p co-stained with miR-674-3p and also cells containing none of the two miRNAs. Data information: Data are shown as number of cells containing tested miRNA and percentage of control using time-matched controls. P value is determined by two-way ANOVA and Tukey’s multiple comparison test. All presented data come from one cohort of animals (n for each group is 6 animals). Significance bars connecting centers of pie charts indicate significance for relative proportion of all 6 subclasses of cells containing miR-155-5p and/or miR-674-3p.

**Figure 7.**
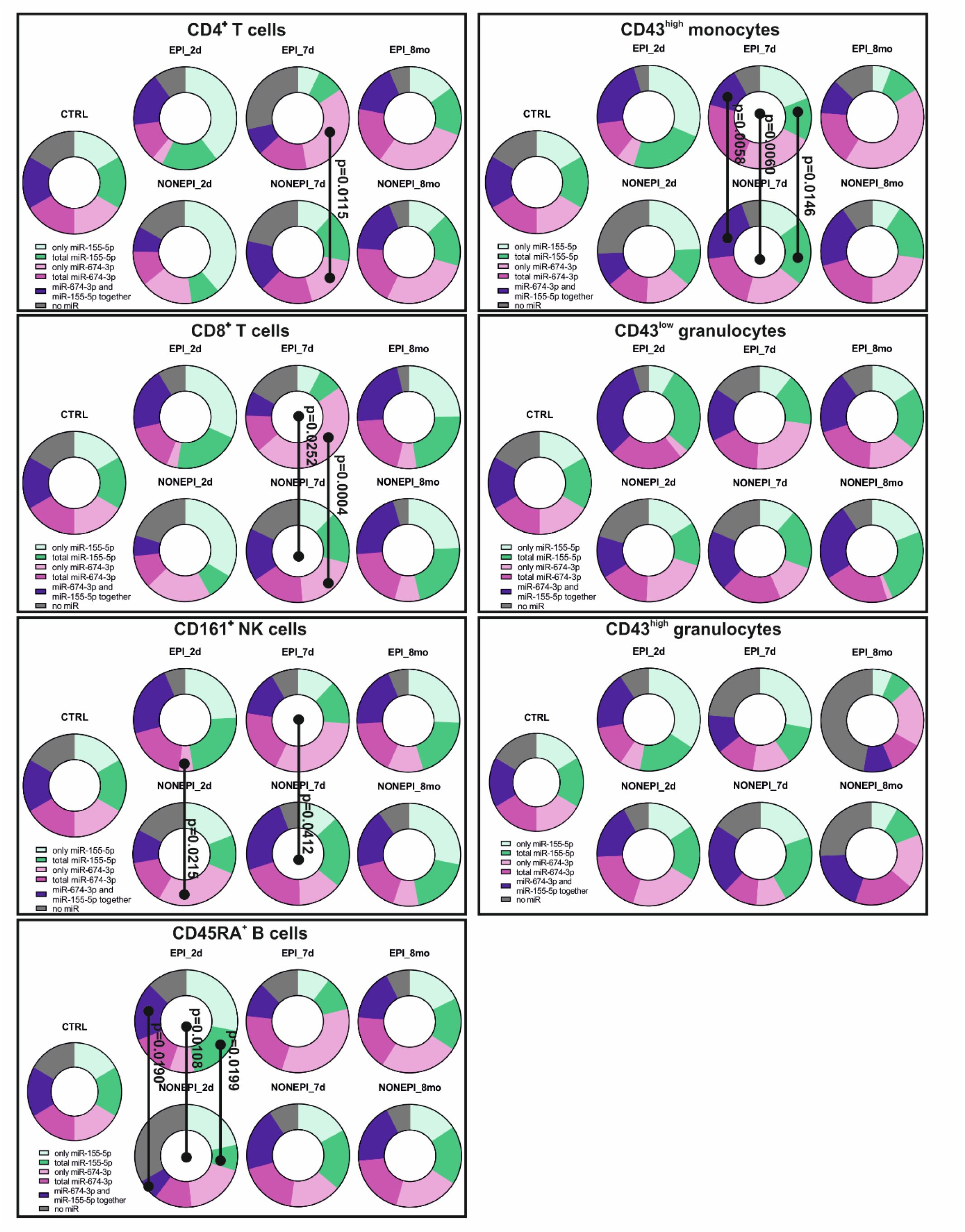
Relative proportion of cell classes containing miR-155-5p and/or miR-674-3p differs between symptomatic and asymptomatic animals at early time points. Pie charts present relative proportion of white blood cells subpopulations containing miR-155-5p and/or miR-674-3p for control, in symptomatic animals that did (EPI) or asymptomatic animals that did not (NONEPI) develop epilepsy, at 2 days, 7 days and 8 months after stimulation. 6 subclasses of cells containing miR-155-5p and/or miR-674-3p: only miR-155-5p (no co-staining with miR-674-3p), total miR-155-5p (could be co-stained with miR-674-3p), only miR-674-3p (no co-staining with miR-155-5p), total miR-674-3p (could be co-stained with miR-155-5p), both miR-155-5p co-stained with miR-674-3p and also cells containing none of the two miRNAs. Data information: Data are shown as number of cells containing tested miRNA and percentage of control using time-matched controls. P value is determined by two-way ANOVA and Tukey’s multiple comparison test. All presented data come from one cohort of animals (CTRL n = 6, EPI n = 3 and NONEPI n=3 animals). Significance bars connecting centers of pie charts indicate significance for relative proportion of all 6 classes of cells containing miR-155-5p and/or miR-674-3p.

**Figure 8.**
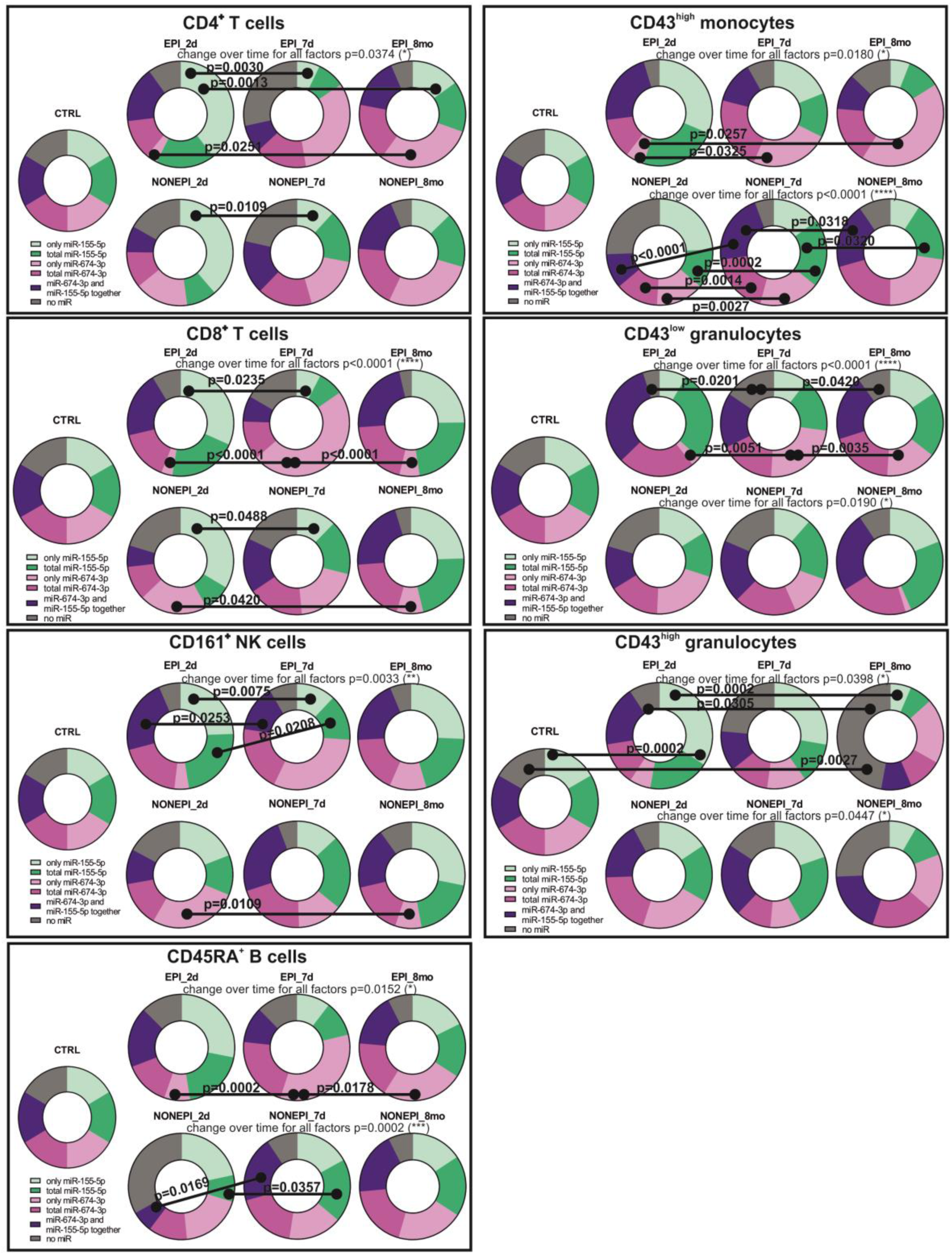
Relative proportion of leukocytes subpopulations containing miR-155-5p and /or miR-674-3p changes in time for symptomatic and asymptomatic animals. Pie charts present relative proportion of white blood cells subpopulations containing miR-155-5p and/or miR-674-3p for control, in symptomatic animals that did (EPI) or asymptomatic animals that did not (NONEPI) develop epilepsy throughout the whole experiment, at 2 days, 7 days and 8 months after stimulation. Control pie charts for each time point look identical as the examples presented on the figure for each cells subpopulation. Each rectangle presents data set for different leukocytes subpopulation. 6 subclasses of cells containing miR-155-5p and/or miR-674-3p: only miR-155-5p (no co-staining with miR-674-3p), total miR-155-5p (could be co-stained with miR-674-3p), only miR-674-3p (no co-staining with miR-155-5p), total miR-674-3p (could be co-stained with miR-155-5p), both miR-155-5p co-stained with miR-674-3p and also cells containing none of the two miRNAs. Data information: Data are shown as number of cells containing tested miRNA and percentage of control using time-matched controls. P value is determined by two-way ANOVA and Tukey’s multiple comparison test. All presented data come from one cohort of animals (CTRL n = 6, EPI n = 3 and NONEPI n=3 animals).

When looking at the selected miRNA presence in white blood cells, we observed a significant difference in the relative proportion of cells containing miR-155-5p and/or miR-674-3p detected between control animals and in rats at 2 days post-stimulation in CD43^high^ monocytes and CD43^high^ granulocytes numbers and at seven days post-stimulation in the number of CD4^+^ T cells, CD45RA^+^ B cells, CD43^high^ monocytes, CD43^low^ granulocytes and CD43^high^ granulocytes (Fig 6 and Suppl Fig 3). There were no significant differences between control animals and stimulated animals at 8 months post-stimulation (Fig 6).

An absolute number of cells containing tested miRNAs, presented as a percentage of control for each time point, are shown on Supplementary Figure 3. At 2 days post-stimulation number of CD43^high^ monocytes containing miR-155-5p and/or miR-674-3p decreased in stimulated animals, when compared to controls, and the number of CD43^high^ granulocytes containing both miRNAs decreased in stimulated animals (Suppl Fig 3A). Additionally, an absolute number of CD4^+^ T cells containing miR-155-5p and/or miR-674-3p decreased in stimulated animals at 7 days post stimulation, and in an absolute number of CD45RA^+^ B cells, CD43^high^ monocytes, CD43^low^ granulocytes, and CD43^high^ granulocytes increased (Suppl Fig 3B).

An absolute number of cells containing one or both miRNAs for all selected subpopulations of white blood cells was also analyzed (Suppl Fig 4). At 2 days post-stimulation, there was a significant increase in the number of cells containing only miR-155-5p, when comparing control and stimulated animals for CD4^+^ T cells (245.1% of control), CD8^+^ T cells (238.1% of control) and CD43^high^ granulocytes (300.3 % of control) (Suppl Fig 4). At 7 days after stimulation, there was an increase in the number of cells containing only miR-674-3p, in subpopulations of CD8^+^ T cells (245.4% of control) and CD45RA^+^ B cells (207.7% of control) (Suppl Fig 4). At 8 months after stimulation, there were no significant changes detected between control and stimulated animals when

### The number of white blood cells subpopulations containing miR-155-5p and/or miR-674-3p differs between asymptomatic and symptomatic animals

We observed there were significant differences between symptomatic animals (EPI group) and asymptomatic animals (NONEPI group) in the relative proportion of cells subpopulations containing miR-155-5p and/or miR-674-3p at 2 or 7 days after stimulation (Fig 7). Contrary, at 8 months post-stimulation there were no significant differences detected between asymptomatic (NONEPI) and symptomatic (EPI) animals (Fig 7).

At 2 days post-stimulation, the relative proportion of 6 CD45RA^+^ B cell classes containing miRNA was significantly changing (p = 0.0108) between EPI and NONEPI animals (Fig 7). Additionally, there was a difference in the relative proportion of CD45RA^+^ B cells containing miR-155-5p (28.09% EPI vs 8.13% NONEPI, p = 0.0199) and relative proportion of CD45RA^+^ B cells containing miR-155-5p and miR-674-3p together (18.35% EPI vs 6.51% NONEPI, p = 0.0190). We also observed a decrease in the relative proportion of CD161^+^ NK cells containing only miR-674-3p in symptomatic animals (4.47%) when compared to asymptomatic animals (27.03%) at 2 days post-stimulation (p = 0.0215).

Significant changes between EPI and NONEPI groups at day 7 post stimulation were detected in subpopulations of cells containing miR-155-5p and/or miR-674-3p presence in CD8^+^ T cells (p = 0.0252), CD161^+^ NK cells (p = 0.0412) and CD43^high^ monocytes (p = 0.060) (Fig 7). Particularly interesting was the increase in the relative proportion of CD4^+^ and CD8^+^ T cells containing only miR-674-3p in symptomatic animals (31.73% and 48.26%, respectively) when compared to asymptomatic animals (17.60% and 19.65% respectively, p = 0.0115 and p = 0.0004) (Fig 7).

In asymptomatic animals (NONEPI group), there was a change over time in the relative proportion of 6 cell subclasses with the presence of miR-155-5p and/or miR-674-3p in CD45RA^+^ B cells (p=0.002), CD43^high^ monocytes (p<0.0001), CD43^low^ granulocytes (p=0.0190) and CD43^high^ granulocytes (p=0.0447) (Fig 8). In symptomatic animals (EPI group), significant differences in the relative proportion of cell subpopulations in time were detected for all tested white blood cells: CD4^+^ T cells (p=0.0374), CD8^+^ T cells (p<0.0001), CD161^+^ NK cells (p=0.0033), CD45RA^+^ B cells (p=0.0152), CD43^high^ granulocytes (p=0.0398), CD43^low^ granulocytes (p<0.0001) and CD43^high^ monocytes (p=0.0180) (Fig 8).

There were also detected differences in the relative proportion of cells containing miRNAs within both symptomatic and asymptomatic animals between all tested time points. Interestingly, there was an increase in the relative proportion of cells containing only miR-155-5p at 2 days after stimulation and then the decrease at 7 days post-stimulation for: CD4^+^ T cells, both in EPI and NONEPI groups (EPI: 39.58% at 2d, 7.09% at 7d, p = 0.0030; NONEPI: 38.7% at 2d, 11.78% at 7d, p = 0.0190), CD8^+^ T cells (EPI: 31.76% at 2d, 7.57% at 7d, p = 0.0235; NONEPI: 33.54% at 2d, 12.31% at 7d, p = 0.0488) and CD161^+^ NK cells in EPI group (24.4% at 2d, 12.03% at 7d, p = 0.0075) (Fig 8). In symptomatic animals, we observed a strong decrease in the relative proportion of cells containing only miR-674-3p at 2 days post-stimulation for: CD4^+^ T cells (3.47%), CD8^+^ T cells (3.38%), CD45RA^+^ B cells (7.86%), CD43^high^ monocytes (5.75%) and CD43^low^ granulocytes (2.80%) (Fig 8). In all these cell types relative proportion of cells containing only miR-674-3p at 7 days post-stimulation was highly increasing: CD4^+^ T cells (31.73%), CD8^+^ T cells (48.26%), CD45RA^+^ B cells (33.92%), CD43^high^ monocytes (23.96%) and CD43^low^ granulocytes (23.68%) (Fig 8). It remained at a higher level at 8 months post stimulation in CD4^+^ T cells (29.48%), CD45RA^+^ B cells (24.74%), CD43^high^ monocytes (42.69%), and CD43^low^ granulocytes (15.38%), decreasing only in CD8^+^ T cells (6.13%) (Fig 8).

Similarly to the relative proportion of cells, absolute numbers of cells containing miR-155-5p and/or miR-674-3p presented as a percentage of control revealed significant change between symptomatic and asymptomatic animals for CD8^+^ T cells (p=0.0252), CD161^+^ NK cells (p=04.12) and CD43^high^ monocytes (p=0.0060) at 7 days post-stimulation (Suppl Fig 5A and B). There were no differences between symptomatic and asymptomatic animals in absolute numbers of cells containing miR-155-5p and/or miR-674-3p at 2 days and 8 months after stimulation.

At 7 days after stimulation, there was a significant increase in absolute numbers of cells between symptomatic and asymptomatic animals in CD4^+^ T (143.87% EPI vs 68.88% NONEPI) cells and CD8^+^ T cells (401.92% EPI vs 88.80% NONEPI) containing only miR-674-3p (Suppl Fig 6A) and the decrease in the absolute number of CD43^high^ monocytes containing miR-155-5p (97.49% EPI vs 293.10% NONEPI) and CD43^high^ monocytes containing both miR-155-5p and miR-674-3p (92.64% EPI vs 303.17% NONEPI) (Suppl Fig 6B).

### The presence of miR-155-5p and/or miR-674-3p in white blood cells allows early prediction of epilepsy

Analysis of 6 cells subclasses revealed a significant increase in the relative proportion of CD45RA^+^ B cells containing miR-155-5p and/or miR-674-3p, presenting an increase in miRNAs levels at 2 days after stimulation (p=0.0108), in animals that would develop seizures at much later time points (Fig 9A). ROC analysis revealed high sensitivity and specificity (AUC=0.7778) for the difference in absolute numbers of CD45RA^+^ B cells containing miR-155-5p and/or miR-674-3p between symptomatic and asymptomatic animals at 2 days post-stimulation (Fig 9B). There was also a decreasing number of CD45RA^+^ B cells containing total miR-155-5p (114.40% EPI vs 29.71% NONEPI) and CD45RA^+^ B cells containing both miR-155-5p and 674-3p (108.3% EPI vs 23.80% NONEPI) at 2 days after stimulation (Fig 9A), making these values strong candidates for predictive biomarkers of early epilepsy development, especially that sensitivity and specificity tested by ROC analysis was 100% (AUC=1.0000) in both cases (Fig 9B). Interestingly in asymptomatic animals, at 2 days post stimulation, there was a significant increase of CD161^+^ NK cells that did not contain neither miR-155-5p, nor miR-674-3p (52.39% EPI vs 154.90% NONEPI) (Fig 9A), ROC analysis revealed 100% sensitivity and specificity (AUC=1.0000) (Fig 9B). Each of these parameters detected at 2 days after stimulation, differentiating between symptomatic and asymptomatic animals, are considered early biomarkers of epileptogenesis in preclinical studies.

**Figure 9.**
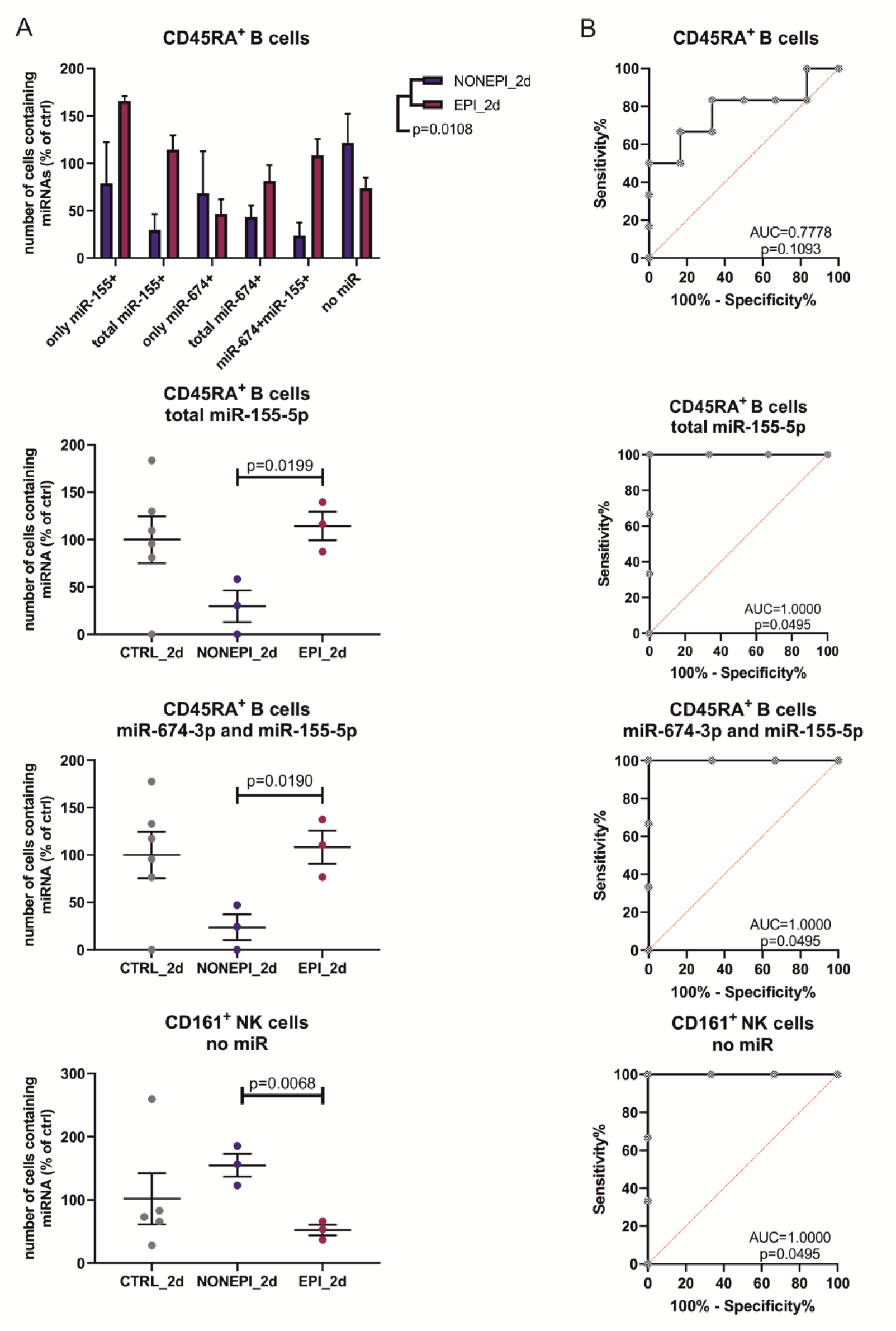
Changes in the number of white blood cells subpopulations, containing miR-155-5p and miR-674-3p, allow predicting at 2 days after stimulation which animals will develop seizures at much later time points. A Graphs present absolute number of cells containing miR-155-5 and/or miR-674-3p at 2 days after stimulation for control, symptomatic animals that did (EPI) or asymptomatic animals that did not (NONEPI) develop epilepsy throughout the whole experiment. B ROC analysis for comparisons of absolute number of white blood cells subpopulations containing miR-155-5p and/or miR-674-3p between EPI and NONEPI animals, for each graphs from panel A. Data information: A Data are shown as mean ± SEM for number of cells containing tested miRNA and percentage of control using time-matched controls. P value is determined by two-way ANOVA and Tukey’s multiple comparison test. B ROC analysis presents % of sensitivity and specificity. Data are shown as area under curve (AUC). All presented data come from one cohort of animals (CTRL n = 6, EPI n = 3 and NONEPI n=3 animals).

In summary, changes in miRNAs signatures in the blood depend on the stimulation of the amygdala, the stage of epilepsy development, and the intensity of epilepsy characterized by seizures number. This first-time, multi-platform, detailed analysis of miRNA level in peripheral blood leukocytes led us to the selection of miR-155-5p and miR-674-3p as potential biomarkers of epileptogenesis. Our data proves that early prediction of epilepsy on the basis of miR-155-5p and/or miR-674-3p presence in white blood cells subpopulations is possible in the preclinical model

## Discussion

The main findings of our study are: (i) miRNA signatures in the whole blood change after epilepsy inducing stimulation; (ii) miRNA signatures in the blood differentiate between presymptomatic and symptomatic animals; (iii) miRNA signatures in epileptic animals differ depending on epilepsy phenotype; (iv) levels of miR-155-5p and miR-674-3p predict development of epilepsy; (v) alterations of miR-155-5p and miR-674-3p levels in the blood following proepileptic stimulus, are cell and time specific; (vi) early changes in number of CD45RA^+^ B cells containing miR-155-5p and/or miR-674-3p are biomarkers of asymptomatic phase of epilepsy; (vii) decrease in number of CD45RA^+^ B cells containing miR-155-5p and miR-155-5p together with miR-674-3p allows early identification of animals that will not develop epilepsy after stimulation; (viii) number of CD161^+^ NK cells containing neither miR-155-5p nor miR-674-3p early after proepileptic stimulus predict development or lack of development of epilepsy later on; (ix) decreased number of CD4^+^ T cells is a biomarker of symptomatic epilepsy.

miRNAs have been tested as potential biomarkers of epileptogenesis and epilepsy in various animal models and in patients suffering from epilepsy as reviewed by: (Henshall, 2014; Henshall *et al*., 2016; Pitkänen *et al*., 2019). Good biomarkers should be easy to detect in clinical or preclinical settings, making easily obtainable blood a promising source of biomarkers (Engel *et al*., 2013; Hanin *et al*, 2020; Henshall, 2014; Lukasiuk & Becker, 2014; Pitkänen *et al*, 2016). Most of the studies searching for the circulating biomarkers of epilepsy are focusing on serum or plasma (Raoof *et al*., 2018; Roncon *et al*., 2015); (Szydlowska *et al*., 2024). We propose that due to well-proven interactions between the immunological system and the brain in neurological diseases, including epilepsy (Vezzani *et al*, 2013a; Vezzani *et al*., 2011; Vezzani *et al*, 2013b), omitting blood cells from analyses may cause exclusion of potentially interesting biomarkers. Thus, in this study, we performed a long-term detailed analysis of miRNA levels in the whole blood and subtypes of white blood cells in order to characterize alterations in leukocytes subpopulations and their miRNA content following proepileptogenic stimulus as well as in the search for biomarkers of epilepsy phenotypes in a preclinical model.

In our longitudinal studies, we observed prolonged changes in miRNAs signature both in control and stimulated animals. Interestingly, our lab previously described similar changes in control animals in the brain (Bot *et al*., 2013). Such alterations could be explained by the maturation and/or aging of animals or by external factors such as seasonal changes (Heegaard *et al*, 2016; Ludwig *et al*, 2019). Changes in miRNA signatures in the blood of control animals over time have important implications not only for understanding the role of miRNAs but also in terms of requirements for designing of experimental studies. In particular there is a need for the use of time-matched controls, especially in long-term experiments. Also, historical samples should not be used in preclinical models mRNA/gene profiling experiments.

We observed global alterations in miRNA signatures in the blood following the epileptogenic stimulus. This is the first dataset from longitudinal studies with several sampling time points. Previously Roncon et al. showed changes in miRNA levels in the plasma of mice with pilocarpine-induced epilepsy at early time points and at 50 days after the first seizure (Roncon *et al*., 2015). Interestingly, Roncon et al. (Roncon *et al*., 2015) showed increased levels of our miRNA of interest, miR-674-3p, in the granule cell layer of the dentate gyrus during the latency phase (4 and 8 days) in the pilocarpine model of epilepsy in mice. However, in this study, Roncon et al. did not use time-matched controls, which makes this result harder to interpret. Brennan et al. also analyzed two time points: early during epileptogenesis and chronic after epilepsy development, however, the latest time point was one month after epilepsy development (Brennan *et al*., 2020). Other studies presenting miRNA level changes in blood focused at only one-time point during symptomatic epilepsy, thus they did not reveal any changes in miRNA signatures during epilepsy development (Liu *et al*., 2010; Raoof *et al*., 2018).

Interestingly in studies involving epileptic patients’ serum or plasma, miRNA testing was done only at one-time point. In some studies, authors analyzed samples 24 hours after a seizure (Brennan *et al*., 2020), but in most of the studies, blood samples were collected at random times during symptomatic epilepsy (Pollard *et al*., 2012; Raoof *et al*., 2018; Wang *et al*., 2015a).

In our study, blood miRNA signatures differentiated between symptomatic and presymptomatic animals, and also depended on epilepsy phenotype in epileptic animals. This is a first study showing that blood parameters could be used as a biomarker of epilepsy development or phenotype in preclinical model.

For further, more detailed evaluation as biomarkers, we selected miR-155-5p and miR-674-3p. To the best of our knowledge, this is the first report indicating that miR-155-5p and miR-674-3p levels are changing peripherally in a time-specific manner during epilepsy development and that they can be used as predictors of epilepsy development or intensity.

It is known that miR-155-5p modulates the interleukin-1 signaling pathway in activated human monocyte-derived dendritic cells (DC) (Ceppi *et al*, 2009). miR-155-5p was already shown to play a role also in epilepsy. Silencing miR-155-5p in rat lithium-pilocarpine model of epilepsy reduces cell apoptosis in the hippocampus by regulating oxidative stress via Sestrin-3 regulation (Huang *et al*, 2018). miR-155-5p regulates inflammatory pathways in pediatric patients with temporal lobe epilepsy and in rat lithium-pilocarpine model of epilepsy. It was also shown that miR-155-5p has a role in the induction of human T cell activation. Analysis of miR-155-5p level in activated T cells revealed a strong increase in its level during the first 24 hours of T cells activation. Authors identified several miR-155-5p targets taking part in transcription co-regulatory activity, basic cellular processes such as endocytosis, cytoskeleton, and subcellular localization, RNA and protein degradation, calcium signaling and modulation of immune cell signaling (Diener *et al*, 2020). The level of miR-155 and pro-inflammatory cytokine TNFα mRNA and protein are upregulated in immature rats subjected to status epilepticus and in LPS-stimulated astrocytes, where the use of TNFα inhibitor downregulated miR-155 expression (Ashhab *et al*, 2013). miR-155 was shown to upregulate TNFα, which in turn promotes the release of inflammatory mediators during the innate immune response in mouse model of LPS/D-galactosamine-induced endotoxin shock (Ceppi *et al*., 2009; Tili *et al*, 2007). miR-155 is in the brain of patients in TLE with hippocampal sclerosis and during epileptogenesis in a rat TLE induced by the electrical stimulation of the hippocampus (Korotkov *et al*, 2018). Experiments performed on human astrocyte cultures stimulated with pro-inflammatory cytokine IL-1β revealed that miR-155 regulates the expression of MMP3, which was previously shown to be elevated in TLE (Gorter *et al*, 2007; Korotkov *et al*., 2018). All these studies clearly indicate the role of miR-155-5p in the inflammation ongoing during epileptogenesis and epilepsy both in patients and in animal models. We do not know what is the relation between miR-155-5p in the brain and the blood, and whether miR-155-5p in our experimental model is produced either in the blood cells or comes from the brain, blood cells, or even other organs. Also, we do not know if it participates in the same molecular pathways in the blood cells and in the brain. This requires further studies. Nevertheless, regardless of its function in the blood of diseased animals, it is a useful biomarker differentiating animals with or without epilepsy and allowing early prediction of which animal becomes epileptic in the future.

We detected miRNA signatures in whole blood, differentiating between animals experiencing high or low number of seizures throughout the experiment. We found that miR-674-3p is a potential biomarker of epilepsy severity in epileptic animals. There is little known about miR-674-3p function in any physiological processes or diseased states. Interestingly, Roncon et al. (Roncon *et al*., 2015) showed increased levels of miR-674-3p in the granule cell layer of the dentate gyrus during the latency phase (4 and 8 days) in the pilocarpine model of epilepsy in mice. Also, in the case of miR-674-3p we do not know what is the source of this miR in the blood cells. We propose, however miR-674-3p as a useful biomarker for preclinical studies.

In the next part of our study, we aimed at the characterization of proposed candidates for biomarkers in individual white blood cell subtypes. We used a newly developed method, PrimeFlow (Lai *et al*, 2018). We could simultaneously identify markers of 7 types of white blood cells and 2 miRs in one run.

We detected miR-155-5p together with miR-674-3p in subsets of main leukocyte subpopulations. In all leukocyte subpopulations we observed cells containing only miR-155-5p, only miR-674-3p, or neither. Interestingly, fractions of cell subpopulation expressing studied miRs were changing in time dependent manner following stimulation. This indicates divergent involvement of cells within leukocyte subpopulations in the disease process.

Interestingly, the number of CD45RA^+^ B cells containing miR-155-5p and/or miR-674-3p at 2 days post-stimulation predicts which animals will or will not develop epilepsy. B cells present antigens and secrete cytokines (Batista & Harwood, 2009). B cells are detected in the diseased brain and can have profound effects (Ortega *et al*, 2020). This is emphasized in multiple sclerosis, in which B cell-depleting therapies are arguably the most efficacious treatment for the condition (Jain & Yong, 2022). In humans, a small number of B cells are rarely found in the parenchyma of the healthy CNS, however, B cells can be recruited through the BBB, meningeal barriers, and choroid plexus (Jain & Yong, 2022). It was also shown that B cells can cross blood-brain barrier in healthy rat brains (Knopf *et al*, 1998). Since B cells were shown to migrate to the injured brain in different neurological diseases, it is possible they also play a role in epileptogenesis/epilepsy development.

We propose that either miR-674-3p level in CD161^+^ NK cells or a relative proportion of B cells containing miR-155-5p and/or miR-674-3p can be used as a predictive tool in epilepsy diagnostics and be of great use in preclinical studies of epilepsy. Additionally, the decreased number of B cells containing total miR-155-5p or B cells containing miR-155-5p together with miR-674-3p allows the identification of animals that will or will not develop seizures after stimulation.

In conclusion, the data presented here provide proof of concept evidence for the use of miRNA level changes in white blood cells as an early biomarker of epileptogenesis.

In addition to miRNA signature changes and cell types containing selected miRNAs, we observed alterations in blood cell numbers. We observed a decrease in the number of CD4^+^ T cells in symptomatic animals. The decreased number of CD4^+^ T cells was previously observed in the immediate postictal state in temporal lobe epilepsy patients. These alterations were more pronounced in patients with hippocampal sclerosis (Bauer *et al*, 2008). Thus, we propose that decreased number of CD4^+^ T cells as a biomarker of symptomatic epilepsy.

Other than the decrease in the number, dysfunctions of T cells were also observed in epilepsy models and human patients. In epilepsy induced by unilateral intrahippocampal kainate injection, Zattoni et al. showed blood-brain barrier disruption and infiltration of T cells and macrophages into the brain (Zattoni *et al*, 2011). Additionally, in mutant mice lacking T- and B-cells, they observed early onset of spontaneous recurrent seizures, suggesting the influence of immune-mediated response on network excitability (Zattoni *et al*., 2011). It was also shown that in the hippocampi of patients suffering from refractory epilepsy, there are strong signs of chronic inflammation and infiltration of different subtypes of lymphocytes, including CD4^+^ T cells (Gales & Prayson, 2017). Thus, it is possible that the decreased number of CD4^+^ T cells in the peripheral blood of epileptic animals in symptomatic epilepsy is caused by increased homing of peripheral immune system cells into the epileptic brain, where they can contribute to epilepsy pathology (Bosco *et al*, 2020; Deprez *et al*, 2011; Gales & Prayson, 2017; Vezzani *et al*., 2011; Zattoni *et al*., 2011). These results, together with our observation on the decreased number of CD4^+^ T cells in the blood in symptomatic epilepsy, suggest that there is a strong interplay between the peripheral leukocytes and epileptic brain. Whether these changes are the result or a cause of further changes in the brain requires further studies.

Our detailed analysis of white blood cell subpopulations containing miR-155-5p and miR-674-3p in control and stimulated animals and in control and symptomatic or asymptomatic animals revealed promising candidates for epileptogenesis/epilepsy biomarkers. Thus, we are proposing 4 predictive biomarkers which allow predicting at 2 days post-stimulation, which animals will or will not develop epilepsy at much later time points. The first proposed biomarker of the asymptomatic phase of epilepsy is a change in the relative proportion of CD45RA^+^ B cells containing miR-155-5p and/or miR-674-3p. The second biomarker is decreased level of total miR-155-5p positive CD45RA^+^ B cells in asymptomatic animals compared to animals with epilepsy. The next biomarker is decreased number of CD45RA^+^ B cells containing both miR-155-5p and miR-674-3p in asymptomatic animals. The last proposed biomarker of an asymptomatic phase of epilepsy is an increased number of CD161^+^NK cells, that do not contain miR-155-5p nor miR-674-3p. Additionally, we found that the decreased number of CD4^+^ T cells allows differentiation of epileptic and non-epileptic animals at 8 months post-stimulation. Thus it can be used as a potential biomarker for identifying epileptic animals in preclinical studies without the need for laborious EEG analysis.

## Methods

### Animal surgery and status epilepticus induction

All animal procedures were approved by the I Local Ethical Committee on Animal Research (permit no. 483/2013, 737/2015 and 889/2019) and conducted in accordance with the guidelines established by The European Council Directives 2010/63/EU.

Adult male Sprague-Dawley rats (270-320 g) from Mossakowski Medical Research Centre Polish Academy of Sciences, in Warsaw (Poland) were used in this study. Rats were housed under controlled conditions (24 °C, humidity 50 - 60 %, 12/12 h light-dark cycle) with food and water available *ad libitum*. Animals were housed in pairs. Environment was enriched by using various toys (wooden or plastic balls, wooden bridges, etc.) and snacks, which were changed every week. Starting four weeks before electrical stimulation, rats were subjected to regular handling every other day for 10 minutes.

The amygdala stimulation model of temporal lobe epilepsy was used in this study as previously described with some modifications (Guzik-Kornacka *et al*, 2011; Nissinen *et al*., 2000). Surgery was performed under isoflurane anesthesia (2-2.5 % in 100 % O2), followed by the injection of butorphanol (Butomidor, Richter Pharma AG, Wells, Austria; 0.5 mg/kg i.p.) for analgesia. A stimulating and recording bipolar wire electrode (Plastic One Inc., Roanoke, VA, # E363-3-2WT-SPC) was implanted into the left lateral nucleus of the amygdala 3.6 mm posterior and 5.0 mm lateral to bregma, 6.5 mm ventral to the surface of the brain. A stainless steel screw electrode (Plastic One Inc., Roanoke, VA, #E363/20) was implanted contralaterally into the skull over the right frontal cortex (3.0 mm anterior and 2.0 mm lateral to bregma) as a surface EEG recording electrode. Two stainless steel screw electrodes were placed bilaterally over the cerebellum (10.0 mm posterior and 2.0 mm lateral to bregma) as grounding and reference electrodes. The contacts of all electrodes were placed in a multi-channel electrode pedestal (Plastic One Inc., Roanoke, VA, #MS363), which was attached to the skull with dental acrylate (Duracryl Plus). After two weeks of recovery, animals were electrically stimulated via the intra-amygdala electrode to evoke status epilepticus. Stimulation consisted of a 100-ms train of 1-ms biphasic square-wave pulses (400 μA peak to peak) delivered at 60 Hz every 0.5 s for 30 min. If the animal did not enter status epilepticus, stimulation was continued for an additional 10 min. The status epilepticus was stopped 1 h after stimulation via an intraperitoneal injection of diazepam (20 mg/kg). If the first dose of diazepam did not suppress status epilepticus, the animal received subsequent doses of diazepam at 5 mg/kg. Time matched control animals had electrodes implanted but did not receive electrical stimulation (sham animals).

Rats were monitored with VIDEO-EEG (Comet EEG, Grass Technologies, West Warwick, RI; Panasonic WV-CP480) continuously from the moment of stimulation until the end of week 5 of the experiment (1^st^ vEEG), from week 11 to week 15 (2^nd^ vEEG) and starting at week 27 until the end of the experiment (3^th^ vEEG). Spontaneous seizures were identified from EEG recordings by browsing the EEG manually on the computer screen. An electrographic seizure was defined as a high frequency (>8 Hz), high amplitude (>2x baseline) discharge lasting for at least 5 s. Latency to the first spontaneous seizure, number and frequency of seizures, and number of epileptic animals in each group were evaluated.

The Discovery cohort was prepared in two tours. The first group of animals consisted of 6 control and 8 stimulated rats, the second group consisted of 5 control and 7 stimulated animals. The discovery cohort consisted of 11 control and 15 stimulated animals. The validation cohort consisted of 5 control and 8 stimulated rats chosen randomly from 4 separate groups of animals. For the flow cytometric analysis, we used blood from a separate cohort of animals that consisted of 6 control and 6 stimulated animals. Only 3 stimulated animals from the PrimeFlow cohort developed seizures. The remaining 3 animals did not develop seizures throughout the whole 8 months of the experiment. It is typical for our animals that after amygdala stimulation, only around half of the animals develops seizures. Experiments for all the cohorts of animals were performed based on identical experimental plan as for the discovery cohort.

### Blood collection

Blood was drawn from anesthetized (brief 3% isoflurane) rats tail vein at 2, 7 days and at 8 months post-stimulation. The blood samples were collected into tubes coated with 1 mg K2EDTA (#363706, BD Vacutainer, USA). All the blood samples were collected between 9 am and 11 am to avoid diurnal miRNA levels fluctuations (Heegaard *et al*., 2016).

For miRNA profiling, samples were snap-frozen using dry ice within minutes after the blood draw and stored at -80°C until use.

For flow cytometry analysis fresh samples were used immediately after the blood draw.

### miRNA isolation and microarray profiling

miRNA was isolated from 500 µl of blood using miRNeasy Kit (#217084, Qiagen). The whole miRNA isolation procedure was performed as described by the kit manufacturer. In the end of the procedure, RNA was eluted to 100 µl of RNAse-free water, aliquoted, and stored at -80°C.

miRNA level was profiled using miRNA 4.1 Array Strip (#902404, Santa Clara, CA, Affymetrix). The procedure was performed according to the manufacturer’s recommendations. 8 µl of RNA was used to perform a poly-A-tailing reaction (AF-902134 GA HWS Kit for miRNA Arrays, Affymetrix). Hybridization was done for 20 hours at +48 °C (AF-900454 GeneChip Hybridization Control Kit, Affymetrix). It was followed by the wash and stain protocol (AF-901910 FlashTagTM Biotin HSR RNA Labeling Kit, Affymetrix). The whole procedure was performed using the GeneAtlas™ instrument from Affymetrix (#00-0393).

### Microarray data analysis

Data analysis of the miRNA 4.1 Array Strips identifying up- and down-regulated miRNAs was performed with RStudio (R v. 3.3.2) using Bioconductor (Gentleman *et al*, 2004), oligo and limma (G, 2005) packages. The microarray data were normalized with the Robust Multi-array Average (RMA) algorithm (oligo package version 1.22.0) (Carvalho & Irizarry, 2010). For the expression analysis, the calculated p-values were based on moderated t-statistics. Furthermore, the Benjamini and Hochberg multiple testing adjustment method was applied to the p-values (FDR – False Discovery Rate). A one-way ANOVA was used to establish miRNA with differential levels between the tested groups.

The correlation, unsupervised clustering, and PCA (principal component analyses) were performed in the RStudio (R version 3.3.2) (R_Core_Team, 2012). PCA was performed on microarray probes using package stats and the lattice package (Sakar & D, 2008). For the heatmap clustering of miRNAs with significantly different levels (p < 0.05), miRNAs were ordered with the clustering complete-linkage method and the Pearson correlation distance measure. The heatmap diagrams were generated with the gplots package (Warnes & GR, 2012). The Pearson correlation test was used to analyze the correlations between miRNAs with significantly different levels (p < 0.05) between stimulated and sham-operated control animals. The fuzzy c-means algorithm implemented in the Mfuzz package (Kumar & E Futschik, 2007) was used to perform clusterization on all probes.

The data discussed in this publication have been deposited in NCBI’s Gene Expression Omnibus (Edgar *et al*, 2002) and are accessible through GEO series accession number GSE234942 (https://www.ncbi.nlm.nih.gov/geo/query/acc.cgi?acc=GSE234942).

### Real-Time PCR

Alterations in levels of selected miRNAs detected with microarrays were validated using the TaqMan system from Applied Biosystems by Life Technologies in a new cohort of animals.

Synthesis of cDNA was performed using TaqMan MicroRNA Reverse Transcription Kit (#4366597, Vilnius, LT, Applied Biosystems by Life Technologies) and nexus gradient Mastercycler (#6331, Eppendorf AG).

The PCR reaction was performed using TaqMan Fast Universal PCR Master Mix (#4367846, Foster City, CA, Applied Biosystems by Life Technologies) and MicroAmp Fast Optical 96-Well reaction plates (#4346906, Applied Biosystems by Life Technologies) in StepOnePlus System (ThermoFisher Scientific). Primers used were: rno-miR-155-5p (RT:002571, ThermoFisher Scientific), and rno-miR-674-3p (RT:001956, ThermoFisher Scientific). Data were analyzed using StepOne Software v. 2.3. Comparative Ct method (ΔΔCt method) was selected for analysis and performed according to ThermoFisher recommendations. As a reference, we used rno-miR-191a-3p (RT:002576, ThermoFisher Scientific), which was selected as the most stable in our experiment, using normfinder algorithm (Andersen *et al*, 2004).

### Phenotyping and miRNA-labelling in peripheral blood cells (PrimeFlow assay)

350 µl of freshly collected peripheral blood was used for analysis at 2 days, 7 days and 8 months after stimulation. Red blood cells were lysed by Ammonium-Chloride-Potassium lysing buffer (Gibco) and incubated on a shaking platform at room temperature till the disappearance of the solution turbidity (∼10 min). White blood cells were collected by centrifugation at 500 g for 5 min and washed twice with PBS with 0,1 % BSA and ice-cold PBS, respectively. Cells were stained with viability dye (FVD eFluor455UV, eBioscience) and the panel of 8 fluorescent antibodies for surface staining (Barnett-Vanes *et al*, 2016): APC-Cy7 for CD45, BV605 for CD3, PE-Cy5 for CD4, BV510 for CD8, BV421 for CD45RA, PE for CD161, FITC for HIS48 and PE-Cy7 for CD43 (see Table 1 for details) in Brilliant Staining buffer (BD Biosciences) for 30 min on ice and washed with Flow Cytometry Staining buffer (eBiosciences). For in situ miRNA hybridization and labeling, the cells were further processed according to an original protocol for PrimeFlow microRNA assay (eBioscience). The target probes for miR-155-5p and miR-674-3p were designed to be recognized by labels with AlexaFluor 647 and 568 fluorochromes, respectively. A minimum of 130,000 viable cells per sample were acquired on a Cytek Aurora flow cytometer (Cytek Biosciences). The cell populations were identified by manual gating, based on FMO controls and strategy to identify the major leukocyte population in rats (Barnett-Vanes *et al*, 2016). Analysis of cellular phenotypes was performed using FlowJo software (BD Life Sciences) and GraphPad Prism v. 8 (Dotmatics) for statistics and visualization. The gating strategy for the PrimeFlow assay is presented in Supplementary Fig 1.

**Table 1.**

| Marker | Target | Fluorophore | Details | Company | Cat.No: |
| --- | --- | --- | --- | --- | --- |
| CD45 | all leukocytes | APC-Cy7 |  | BioLegend | 202216 |
| CD3 | all lymphocytes | BV605 |  | BD | 563949 |
| CD4 | T helper cells | PE-Cy5 |  | BD | 554839 |
| CD8 | T cytotoxic cells | BV510 |  | BD | 740139 |
| CD45RA (B220; HIS24) | B cells | V450 |  | BD | 561626 |
| CD161 | NK / NKT cells | PE |  | BioLegend | 205604 |
| HIS48 | Granulocytes | FITC |  | eBioscience | 11-0570-82 |
| CD43 | Neutrophils | PE-Cy7 |  | BioLegend | 202816 |

### Statistics

The statistical analysis of microarray data, Real-Time data, and Prime Flow data was performed using Graph Pad Prism (v.8, GraphPad Software, USA). Two-way ANOVA with Tukey’s multiple comparisons test, Sidak’s multiple comparison test, or unpaired t-test were used. The statistical significance was determined as p < 0.05.

We confirm that we have read the Journal’s position on issues involved in ethical publication and affirm that this report is consistent with those guidelines.

## Author contributions

Conceptualization: KS and KL. Methodology: KS, PC. Investigation: KS, PC, KN, MO. Data curation and Formal Analysis: KS and PC. Visualization: KS and PC. Writing–original draft preparation: KS. Writing—review and editing: KS, PC and KL. Supervision: KL and KP. Funding Acquisition: KL. All authors have read and agreed to the published version of the manuscript.

## Data Availability

The data discussed in this publication have been deposited in NCBI’s Gene Expression Omnibus (Edgar et al, 2002) and are accessible through GEO series accession number GSE234942 (https://www.ncbi.nlm.nih.gov/geo/query/acc.cgi?acc=GSE234942).

## Ethics approval and consent to participate

All animal procedures were approved by the I^st^ Local Ethical Committee on Animal Research in Warsaw (permit no. 483/2013, 737/2015, and 889/2019) and conducted in accordance with the guidelines established by The European Council Directives 2010/63/EU.

## Consent for publication

Not applicable.

## Acknowledgements

We would like to acknowledge Aleksandra Stepniak for her help with the plasma samples collection and occasional help with the animal care.

