## Supplementary figures and images for "miR-155-5p/miR-674-3p presence in peripheral blood leukocytes and relative proportion of white blood cell types as biomarkers of asymptomatic and symptomatic phases of temporal lobe epilepsy"

### Supplemental Figure 1

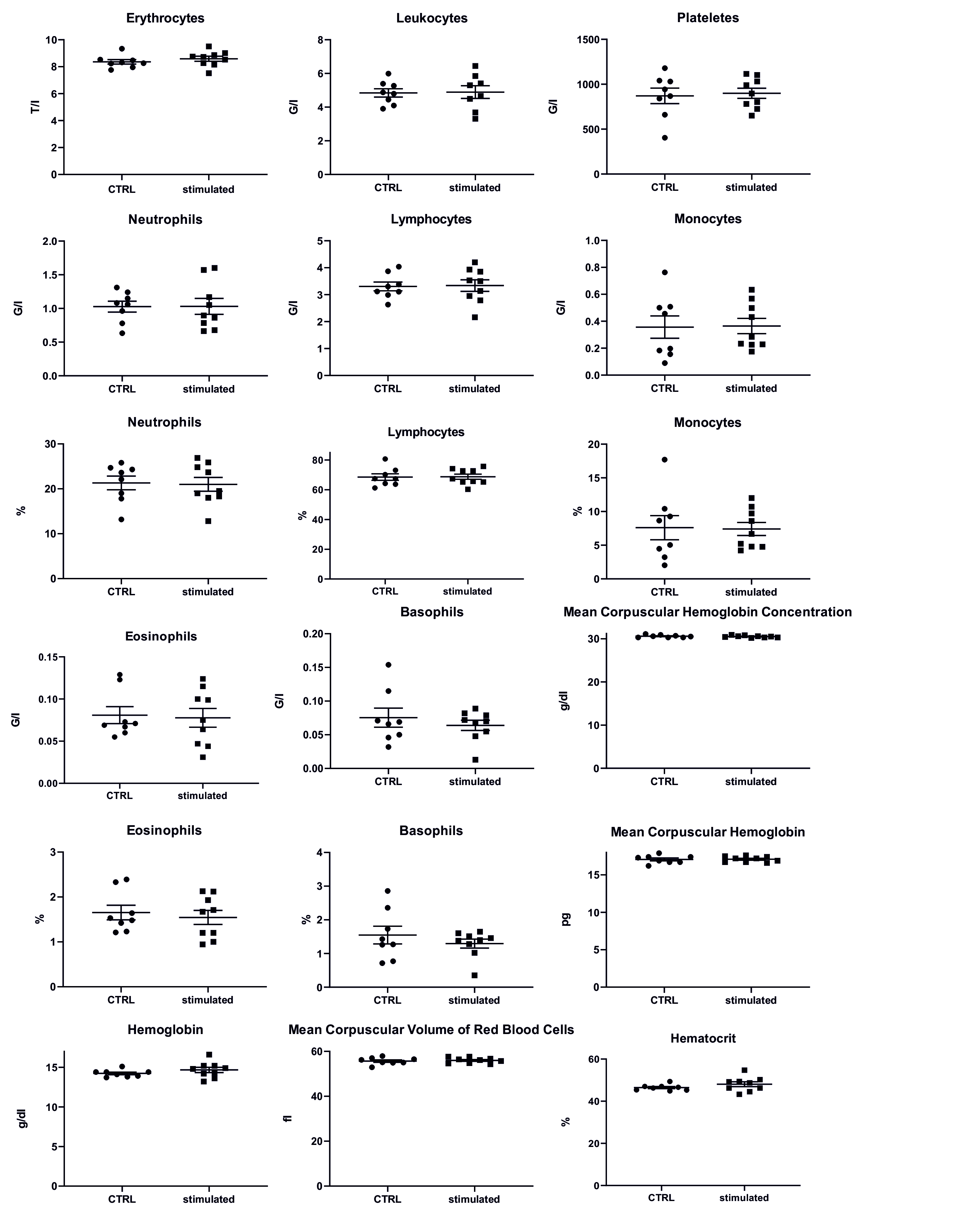

### Supplemental Figure 2

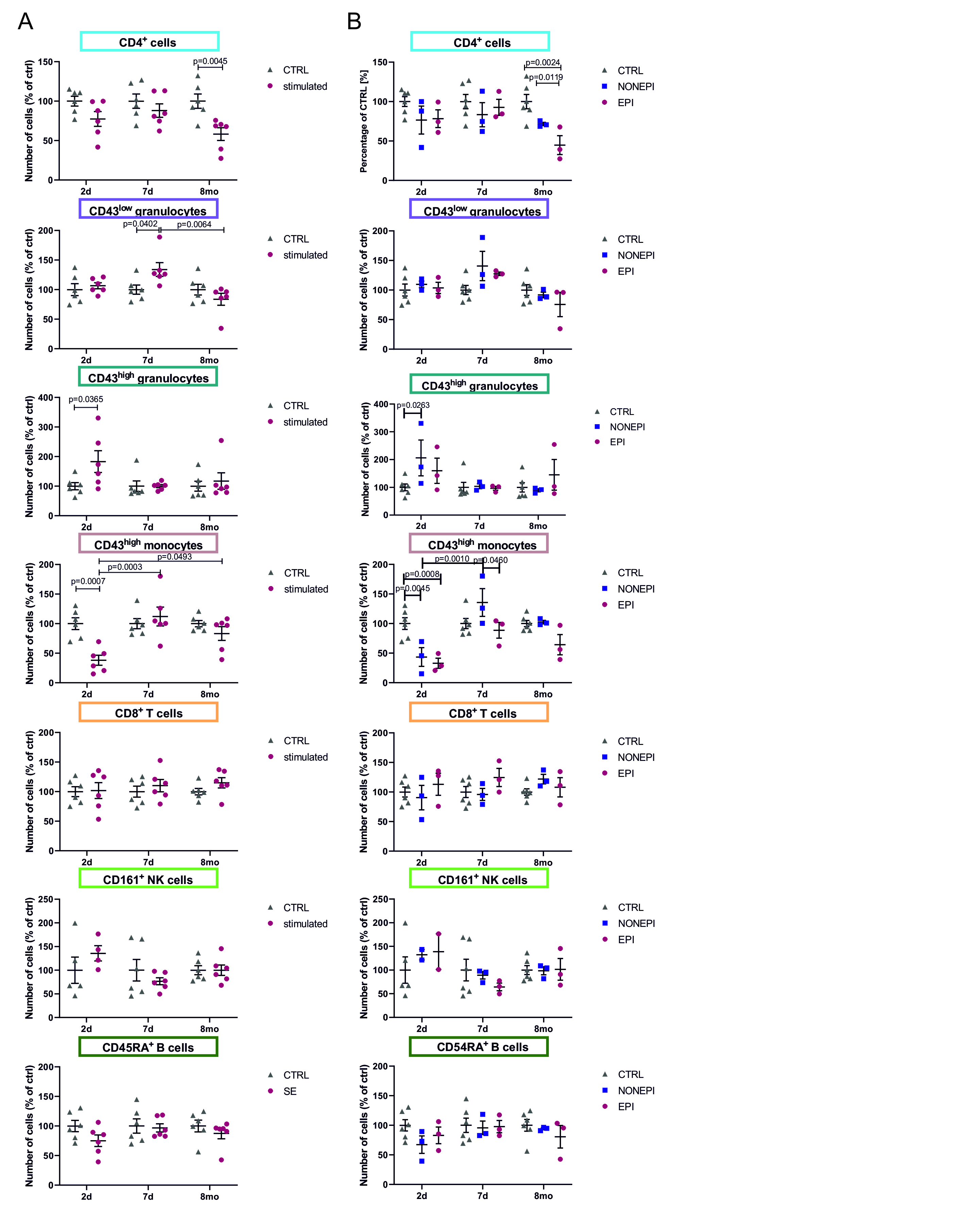

### Supplemental Figure 3

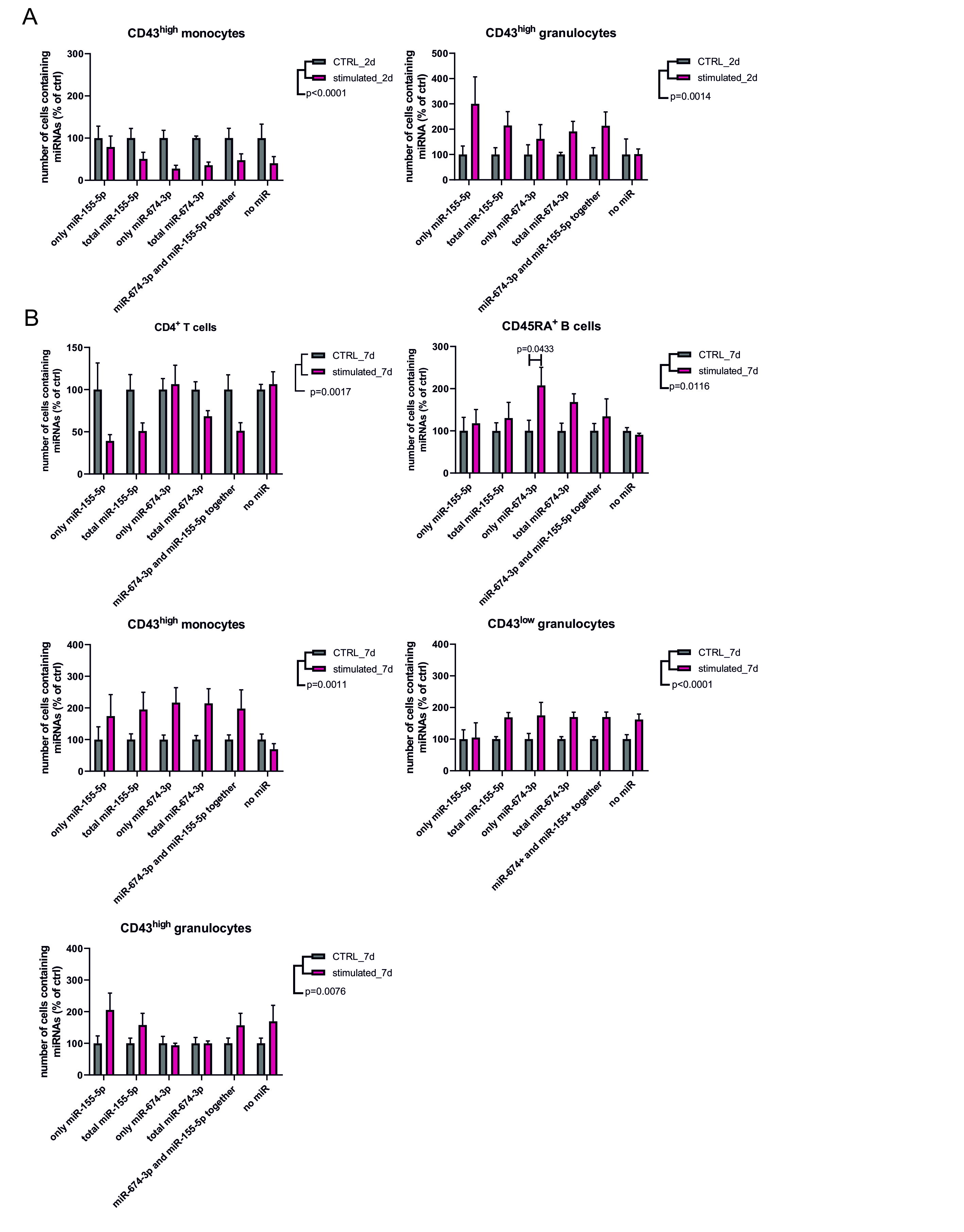

### Supplemental Figure 4

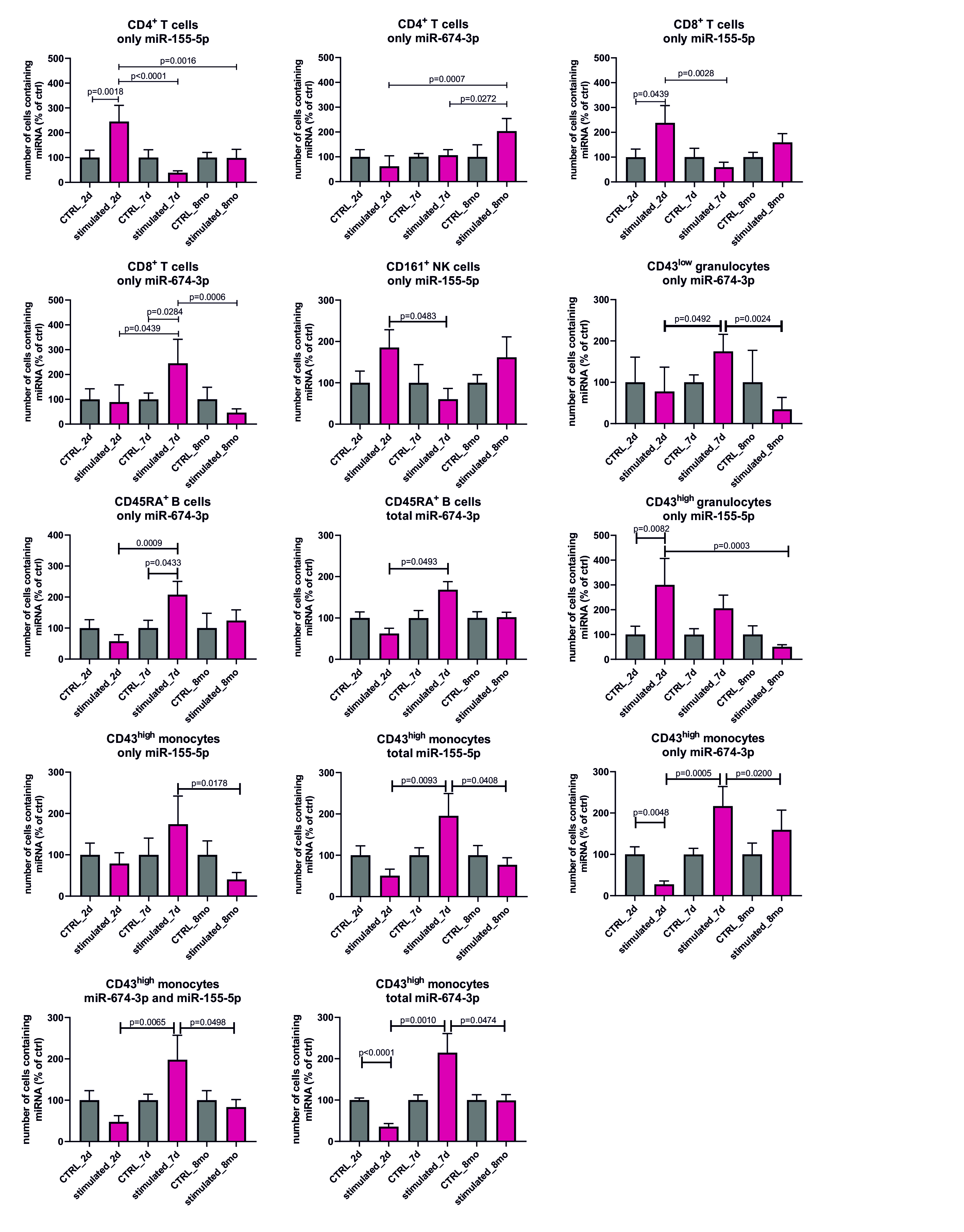

### Supplemental Figure 5A

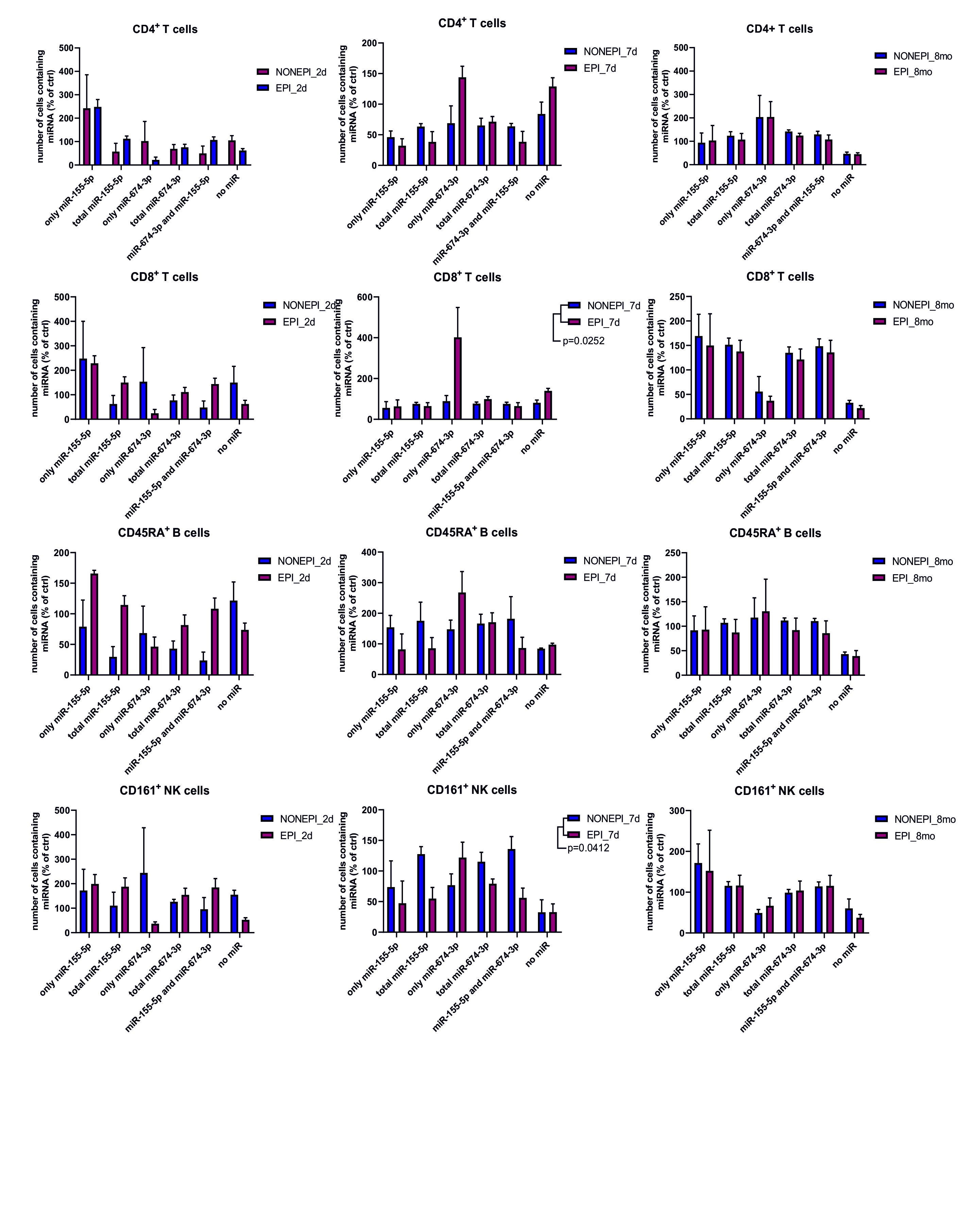

### Supplemental Figure 5B

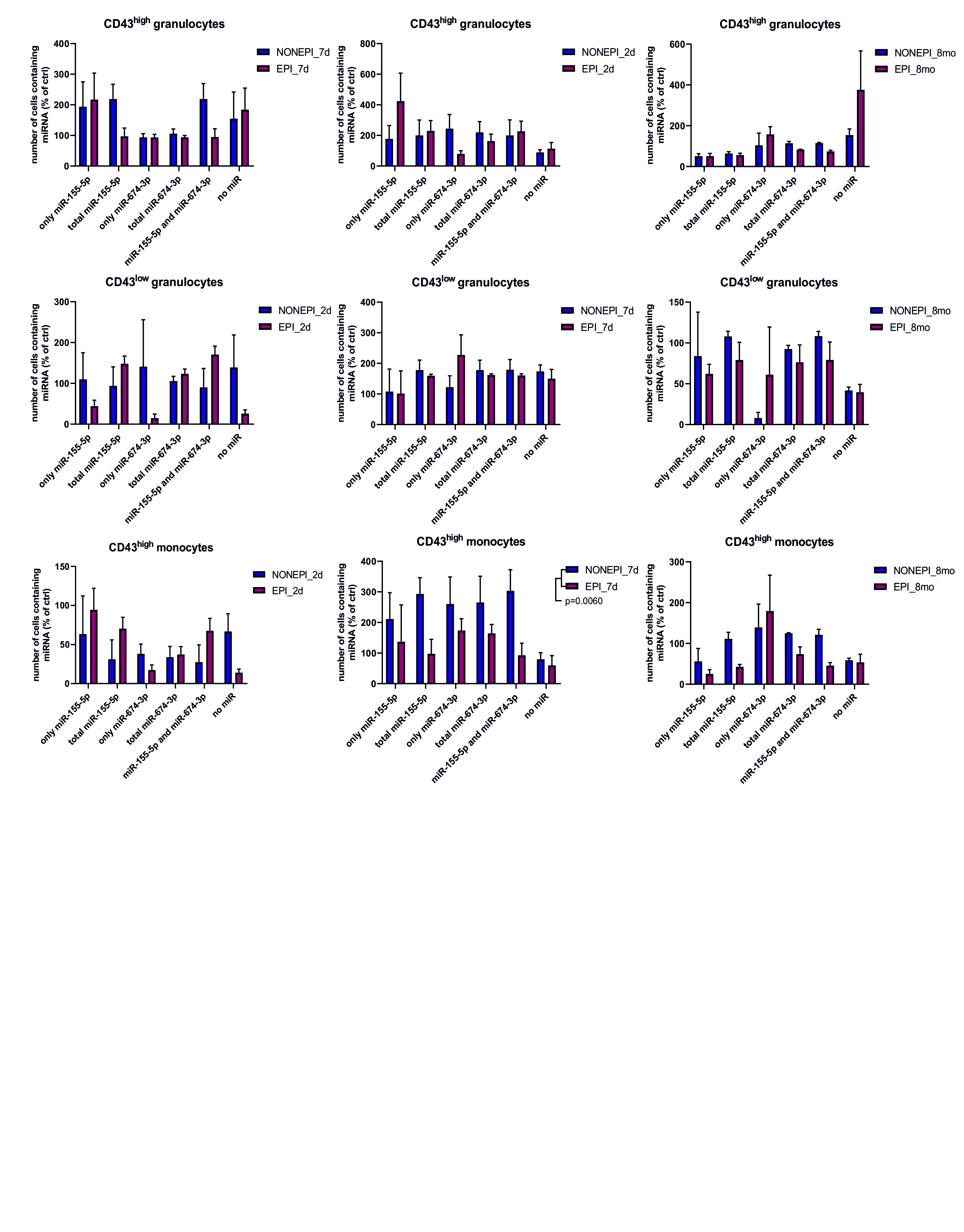

### Supplemental Figure 6A

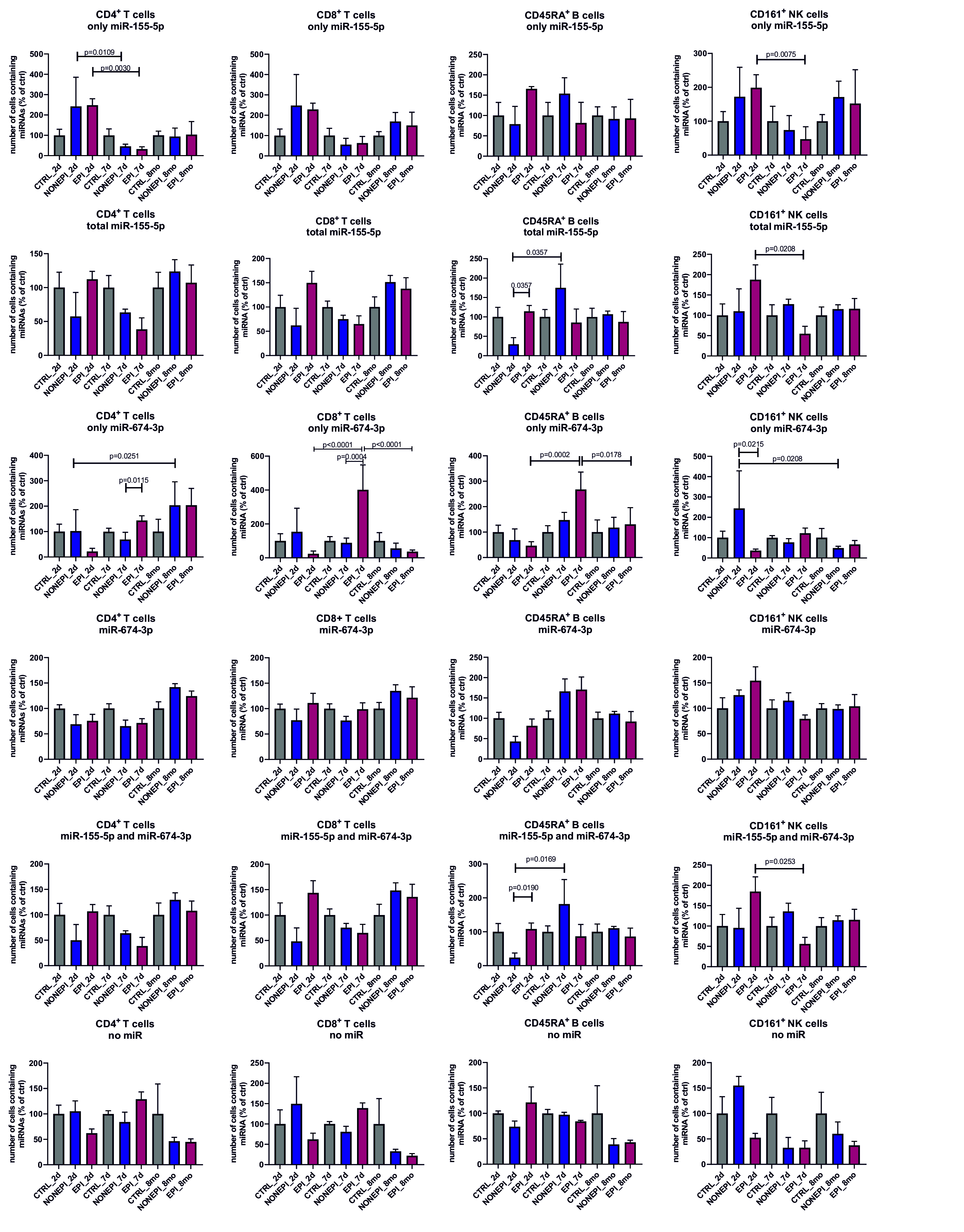

### Supplemental Figure 6B

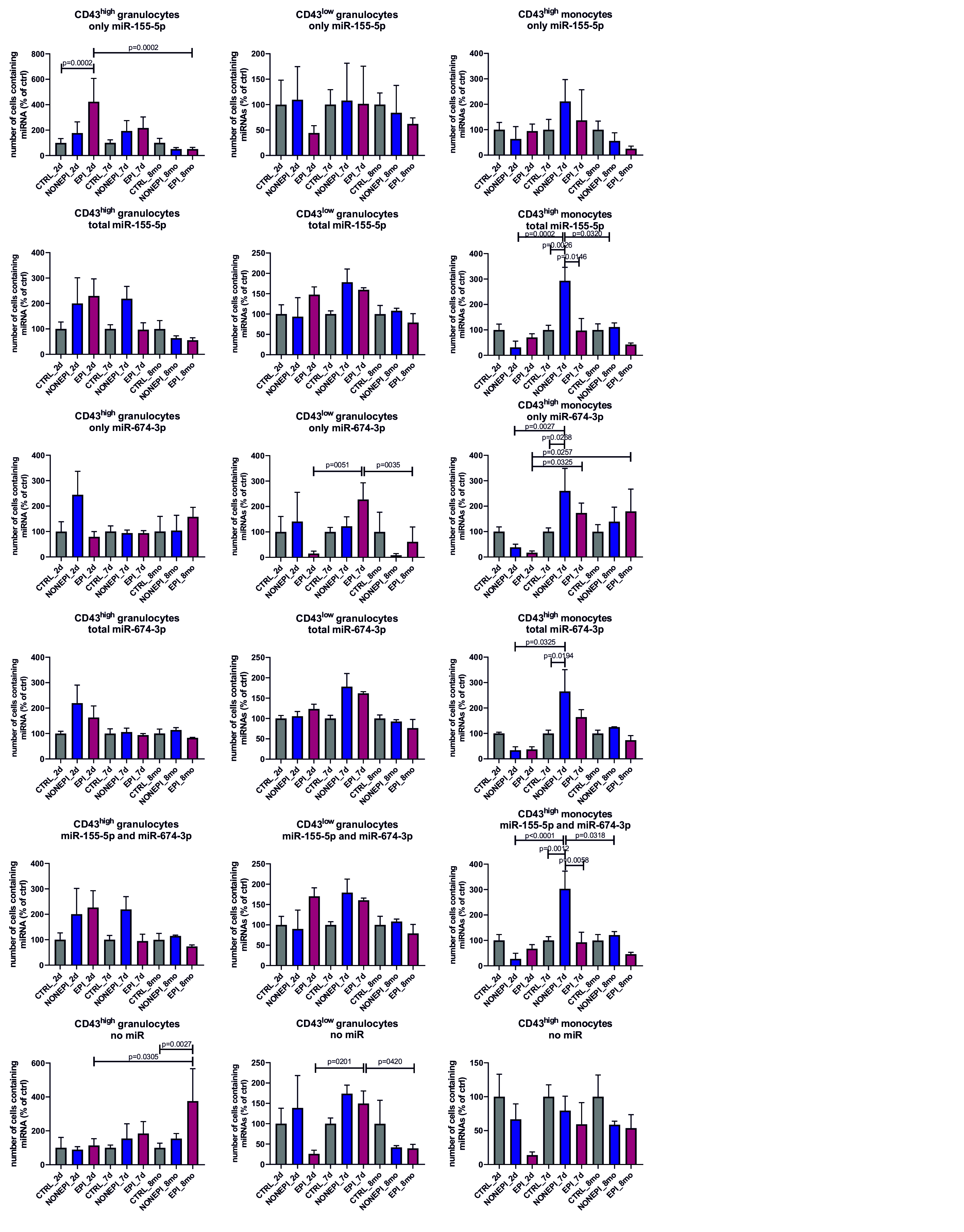

### Supplemental Figure 7

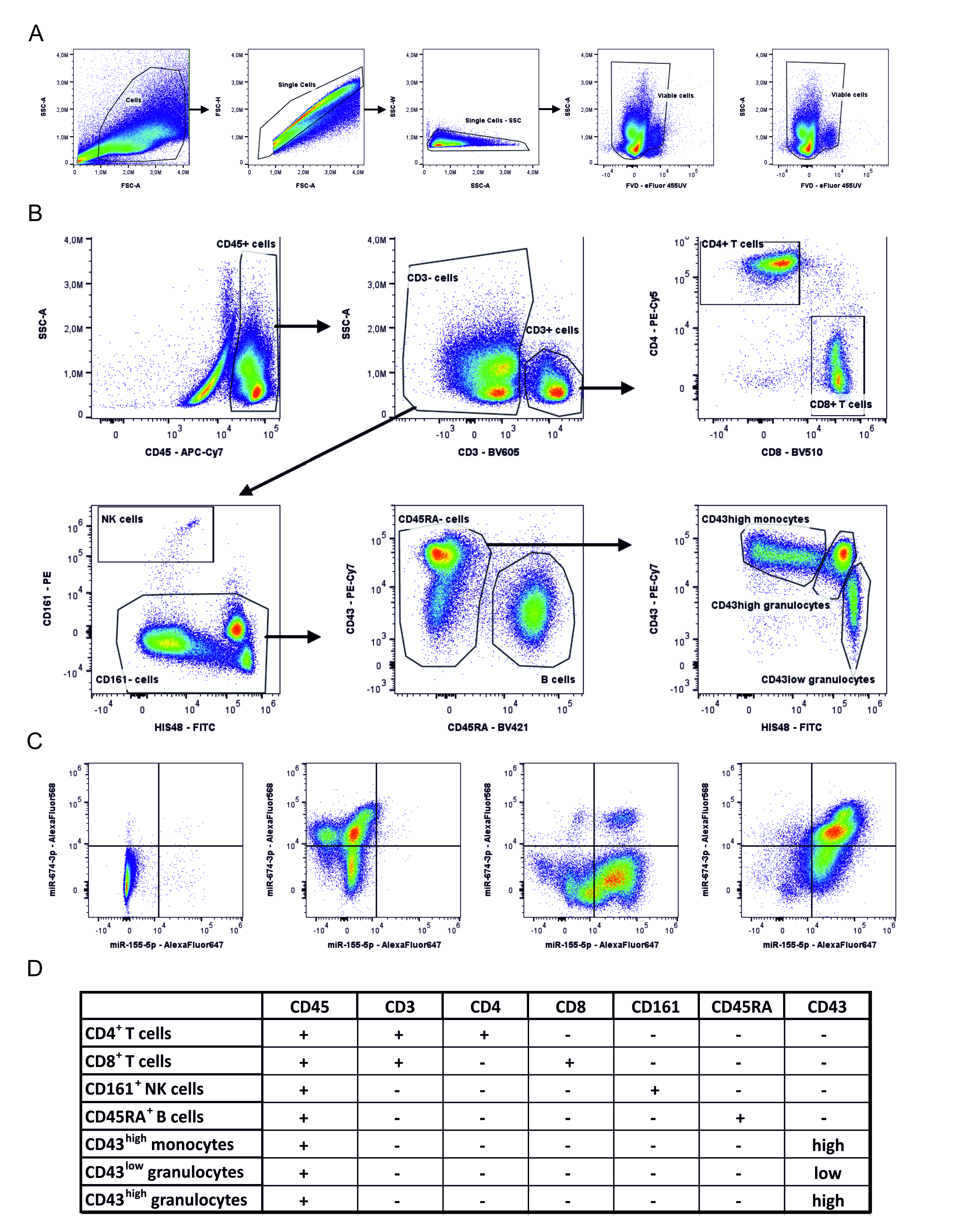
